# Scalable assessment of genome editing off-targets associated with genetic variants

**DOI:** 10.1101/2024.07.24.605019

**Authors:** Jiecong Lin, My Anh Nguyen, Linda Y. Lin, Jing Zeng, Archana Verma, Nola R. Neri, Lucas Ferreira da Silva, Adele Mucci, Scot Wolfe, Kit L Shaw, Kendell Clement, Christian Brendel, Luca Pinello, Danilo Pellin, Daniel E. Bauer

## Abstract

Genome editing with RNA-guided DNA binding factors carries risk of off-target editing at homologous sequences. Genetic variants may introduce sequence changes that increase homology to a genome editing target, thereby increasing risk of off-target editing. Conventional methods to verify candidate off-targets rely on access to cells with genomic DNA carrying these sequences. However, for candidate off-targets associated with genetic variants, appropriate cells for experimental verification may not be available. Here we develop a method, Assessment By Stand-in Off-target LentiViral Ensemble with sequencing (ABSOLVE-seq), to integrate a set of candidate off-target sequences along with unique molecular identifiers (UMIs) in genomes of primary cells followed by clinically relevant gene editor delivery. Gene editing of dozens of candidate off-target sequences may be evaluated in a single experiment with high sensitivity, precision, and power. We provide an open-source pipeline to analyze sequencing data. This approach enables experimental assessment of the influence of human genetic diversity on specificity evaluation during gene editing therapy development.

## INTRODUCTION

In the rapidly advancing field of therapeutic genome editing, ensuring comprehensive characterization of the specificity of genome modification technologies is crucial to developing well-characterized and safe therapies^1,2^. A significant challenge for these technologies is posed by off-target editing, where sequence-specific editing tools such as engineered forms of RNA-guided Clustered Regularly Interspaced Short Palindromic Repeats (CRISPR) systems, may bind and introduce modifications to sequences elsewhere in the genome which are similar to the intended target^3–5^. These modifications can include substitutions, short indels, and structural variants, including translocations between the on-target and off-target locus. Off-target effects could have an impact on the functional potential of edited cells, but off-target edits are challenging to predict and control^6^. The number and type of mismatches or bulges between the guide RNA (gRNA) and target sequence are only poorly correlated with the likelihood of cleavage. Models that use free energy parameters and/or deep learning have modest predictive power, but the likelihood of a candidate off-target being verifiable as an actual off-target in cells remains difficult to predict, as it may depend on the nature of editor delivery to cells and be impacted by the concentration of and duration of exposure to the editor as well as the cellular genomic context^7^. Genetic variation further complicates this issue by producing allele-specific off-target sites^8,9^. We and others have developed software to nominate candidate variant-aware off-target sites using genetic variant datasets as input^9–11^. We found that a genetic variant common in African ancestry populations introduces an experimentally verifiable off-target site for a gRNA targeting the *BCL11A* +58 erythroid enhancer that is part of the recently FDA-approved Cas9 gene edited cell therapy for sickle cell disease and β-thalassemia (exa-cel)^10^. This experimental verification relied on serendipitous access to primary hematopoietic stem and progenitor cells (HSPCs) from an individual carrying this variant that could be subject to gene editing and off-target genotyping. However, the vast majority of genetic variants associated with candidate off-target sites are rare and access to primary cells carrying relevant genetic variants for experimental verification of candidate off-targets is typically infeasible. This problem extends to genetic variants across ancestral distribution and allele frequency as primary cells such as CD34+ HSPCs are relatively scarce and donors do not typically undergo whole genome sequencing prior to acquisition.

Regulatory guidance has recognized the challenge of genetic variant associated off-target potential, requesting investigators to consider the impact of genetic variation on genome editing specificity^12^. For exa-cel, the FDA required a post-marketing surveillance study to screen subjects for candidate genetic variants associated with off-target editing potential and to evaluate for off-target editing in clinical samples^13^. The package inserts states: “Off-Target Genome Editing Risk: Although not observed in healthy donors and patients, the risk of unintended, off-target editing in CD34+ cells due to uncommon genetic variants cannot be ruled out”^14^, indicating that this topic remains an area of uncertainty regarding the safety and specificity of gene editing therapies.

As the field progresses towards the development of new tools and the number of clinical applications increases, the need for accurate and comprehensive quantitative methods for better understanding how donor variability, editing protocol, delivery systems, gRNA design, and cellular state impact off-target editing is crucial. This could improve the pre-clinical evaluation of genetic variants associated with off-target editing potential and could support the clinical implementation of therapeutic genome editing. Furthermore, quantifying off-target editing from multiplexed assay data is a challenging problem requiring estimating low-frequency events in heterogeneous genomic contexts with varying background noise levels.

In scientific literature, the concept of inserting specific sequences of interest into the genomes of target cells using lentiviral vectors has emerged as a powerful tool for characterizing the activity of genome editing tools^15–18^. In a recent study, authors propose using base editors to introduce variants of uncertain significance and investigate their functional consequences in tumorigenesis and drug sensitivity^19^. However, defining effective combinations of gRNAs and base editors for the high-throughput testing of cancer-associated single nucleotide variants remains challenging. To achieve this goal, (Sánchez-Rivera et al., 2022) developed a modular ‘base editor sensor’ platform that couples a gRNA with its cognate genomic target within a lentiviral vector. This design enables measurement of editing efficiency and precision of thousands of sgRNAs paired with functionally distinct base editors, providing a comprehensive resource for introducing and interrogating the effect of cancer-associated variants. The SURRO-seq^20^ method builds upon the results of the CRISPRon method^21^, developed by the same research group. Specifically designed for evaluating the editing activity of RNA-guided nucleases, SURRO-seq involves transducing SpCas9-overexpressing HEK293T cells using a pool of lentiviral surrogate vectors carrying surrogate off-target sequences. To enhance the precision of estimating editing activity, the authors employ a barcode strategy that allows demultiplexing of sequencing files based on the tested off-target sequence. However, these existing methods have several limitations. None leverage Unique Molecular Identifiers (UMIs) for accurate quantification. The use of cell lines overexpressing Cas9 may not faithfully reflect the clinical editing protocol and therefore provide inaccurate estimations due to differing conditions. Furthermore, existing methods show relatively high background noise levels, close to that of clinically relevant off-target editing, thus limiting sensitivity to detect off-targets.

From a computational perspective, several bioinformatics analysis tools have been developed for estimating genome-editing activity rates from experimental data. Methods such as CRISPResso2^22^, ampliCan^23^, and CRISPECTOR^24^ allow for the detection of lower-level activity by comparing indel frequencies using treated vs untreated approach. CRISPResso2 and ampliCan’s main limitation is the lack of a rigorous inferential procedure for estimating editing confidence intervals and statistical testing for indel enrichment. CRISPECTOR uses a two-step approach consisting of a Bayesian classifier for discriminating indels caused by editing activity from background noise, and the estimation of indel proportion using a normal standard approximation. However, this approximation may be unreliable for probabilities close to either 0 or 1, as with off-target and on-target editing. In addition, the CRISPECTOR framework does not allow the inference of editing activity from data generated using more complex experimental designs, including, for example, multiple technical and or biological replicates and donors.

Here we introduce ABSOLVE-seq (Assessment By Stand-in Off-target LentiViral Ensemble with sequencing), a comprehensive workflow that combines a wet lab protocol and an analytical pipeline for estimating editing rates at a set of potential off-target sequences associated with genetic variants. Our approach offers several advantageous features. It has the ability to deliver a pool of candidate sequences as a lentiviral library, allowing scalable multiplexed testing of numerous candidate off-target sequences in a single experiment. The library can be delivered to primary cells which in turn can be edited by a clinically relevant gene editor delivery method^25^. It includes UMIs and candidate barcodes in the vector sequence and combines information from plasmid pool sequencing and experimental samples. This setup offers the possibility to analyze data using different quantification metrics and improves the assay’s precision and sensitivity by suppressing oligo synthesis, packaging, amplification, and sequencing errors. It also can test reference and alternative alleles for the same candidate off-target site in parallel to investigate the allele-specific nature of editing. The ABSOLVE-seq analytical pipeline builds on the CRISPResso framework, extending and improving the quantification and estimation of indel frequency using a logistic regression model.

Our method estimates the difference in indel frequency between edited and unedited samples (Δ indels) at each candidate off-target, calculates marginal distributions and corresponding confidence intervals, and performs statistical testing. Using a generative model calibrated on the experimental data for each candidate off-target, we calculate the statistical power of detecting significant positive Δ indels for a given indel frequency threshold.

For experimental validation, we employ two *BCL11A* enhancer-targeting guide RNAs currently in clinical use or development for sickle cell disease and β-thalassemia^25–28^. As genome editing moves into the clinic, ABSOLVE-seq can support quantitative assessment of editing at candidate off-target sequences involving genetic variants, enabling a more comprehensive population-level risk analysis for new therapies.

## RESULTS

### The ABSOLVE-seq framework

We introduce ABSOLVE-seq, a comprehensive workflow that combines a wet lab protocol and an analytical pipeline to estimate editing frequency for a set of candidate variant-associated off-target sequences in a single multiplexed experiment.

The workflow begins with nominating a set of candidate off-target sites for a gRNA sequence. Here we focus on the use case of genetic variant-associated off-target sites since these are typically infeasible to test in a relevant cellular context given cells carrying the endogenous variants are not typically available. We use the CRISPRme^10^ tool to nominate candidate variant-associated off-target sites (**Figure 1A**). Subsequently, a library of lentiviral vector constructs is synthesized (**Figure 1B**), with each construct carrying a single candidate off-target sequence. We add a candidate-specific 9-bp barcode positioned away from the predicted cleavage site to minimize risk of edits disrupting the barcode, which enables candidate identification even in the presence of editing. Following PCR library construction, each candidate-barcode pair is linked to numerous 16-bp UMIs (**Supplementary Figure 1)**. The library is then packaged in lentiviral vectors. After transduction of cells by the lentiviral pool, the sample undergoes the therapeutic gene editing protocol, allowing for parallel testing of numerous candidate off-target sequences, as well as controls, in a single experiment. DNA is then isolated from edited cells as well as unedited cells cultured in parallel. Sequencing libraries are prepared from edited DNA, unedited DNA, as well as the initial plasmid pool (**Figure 1C**). Distinct samples can compare results across different donor cells, cell states, gene editors, and editor delivery conditions.

**Figure 1.**
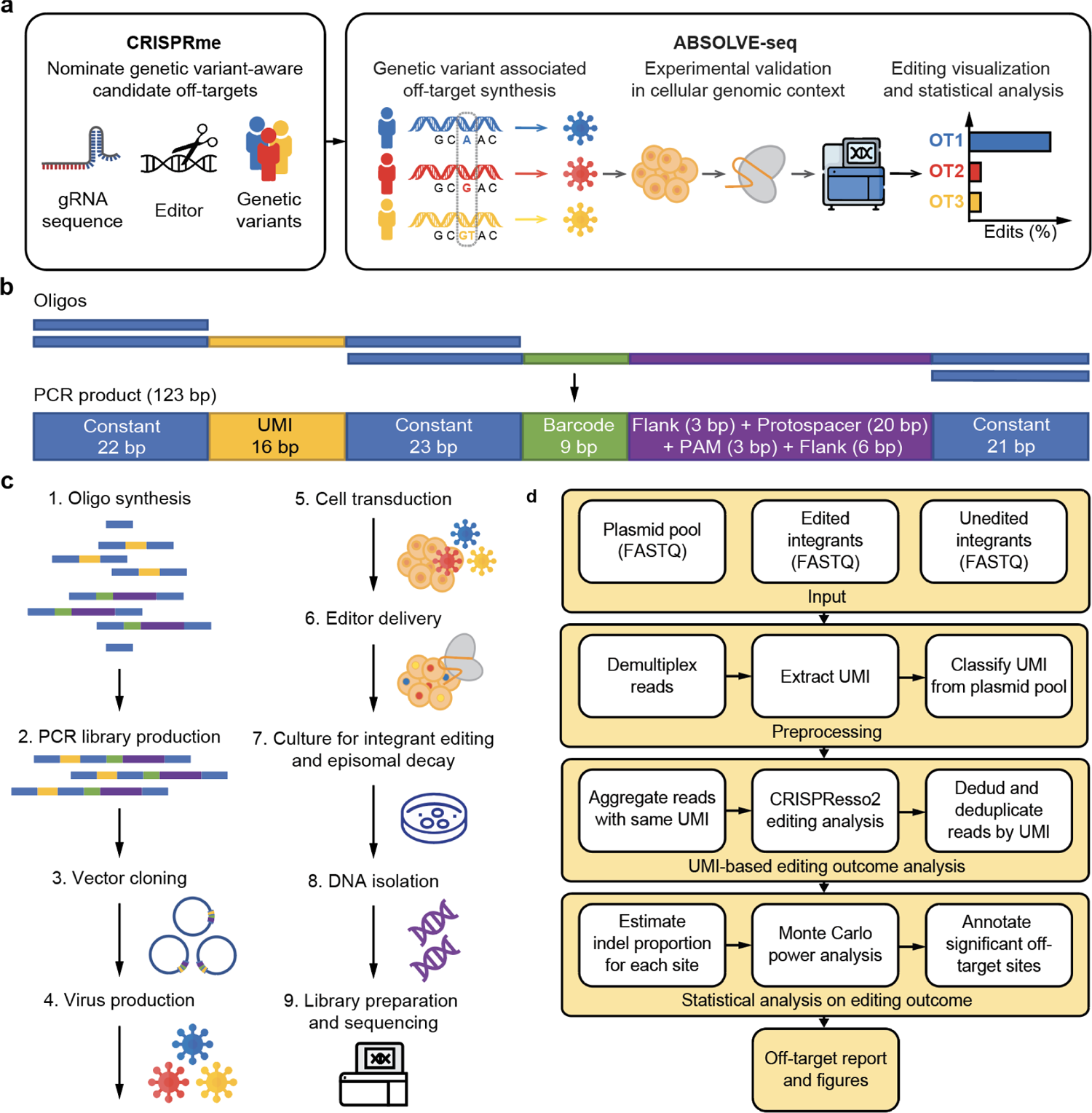
Overview of the ABSOLVE-seq framework. (A) Proposed strategy for assessing genome editing specificity with consideration for human genetic diversity. (B) Schematic of the ABSOLVE-seq oligo design, including a candidate off-target sequence of interest (protospacer + PAM with flanking endogenous sequence), barcode identifying the candidate, and UMI identifying the parent molecule. (C) Overview of the ABSOLVE-seq experimental protocol. (D) Overview of the ABSOLVE-seq computational analysis pipeline.

The sequencing data, along with a table reporting sample annotations, are then provided as input to the ABSOLVE-seq computational pipeline (**Figure 1D**). The raw input data are combined for preprocessing, and we generate candidate-specific sequencing files for analysis using CRISPResso2^22^. The output is further analyzed using various strategies to quantify coverage and indel frequency at the candidate sequences. The data from these analyses, collected from multiple samples, is then consolidated into a single dataset for subsequent statistical modeling.

The indel frequency data are modeled using a logistic regression model (for binomial data) to estimate Δindels. Relevant sample annotations are included in the model as covariates, allowing for consideration of potential confounding factors such as heterogeneous sequencing depth and technical and donor variability. The Δ indels marginal prediction and confidence interval are calculated, and statistical testing is performed. The statistical power of detecting different effect sizes is evaluated using a Monte Carlo procedure, with a generative model calibrated on the data from the unedited samples for each candidate variant-associated off-target.

### Integrant sequences provide reliable estimates for different levels of editing

We first tested the ability of the ABSOLVE-seq integrant structure to accurately report editing of on-target sequences. We tested the gRNA sg1617^25^ used in exa-cel, the first FDA-approved gene edited cell therapy, which disrupts the *BCL11A* +58 erythroid enhancer and induces fetal hemoglobin (HbF) for β-hemoglobinopathy patients. We also included a second gRNA targeting the *BCL11A* +55 enhancer (sg1450), which is also a therapeutic candidate for gene editing to induce HbF for β-hemoglobinopathies^28^. We cloned the *BCL11A* enhancer +58 and +55 on-target sequences to the barcoded and UMI-tagged ABSOLVE-seq vector. CD34+ HSPCs from 3 donors were transduced with lentiviral vectors carrying the integrant on-target sequences. 16 hours later, they were electroporated with 3xNLS-SpCas9-HiFi:sg1617:sg1450 ribonucleoprotein (RNP). After *in vitro* cell culture for an additional 7 days to allow for editing to occur and for decay of episomal lentiviral genomes, we observed vector copy number of 5.2 vector copies per diploid cell (**Supplementary Figure 2**). To quantify editing, genomic DNA was used to prepare next generation sequencing libraries to evaluate indel frequency. The endogenous and lentiviral integrant sequences from the same cells were amplified, sequenced, and analyzed in parallel. We observed similar editing frequencies for the endogenous and integrant sequences for both on-targets (**Figure 2A**). To evaluate integrant sequence quantification across a range of editing frequencies, we compared endogenous and integrant indel frequencies for sg1450 with RNP concentrations ranging from 0.1 to 5 µM. We observed similar editing results for the endogenous and integrant sequences across a wide range of indel frequencies, from approximately 0.1% to 100% indels (**Figure 2B**).

**Figure 2.**
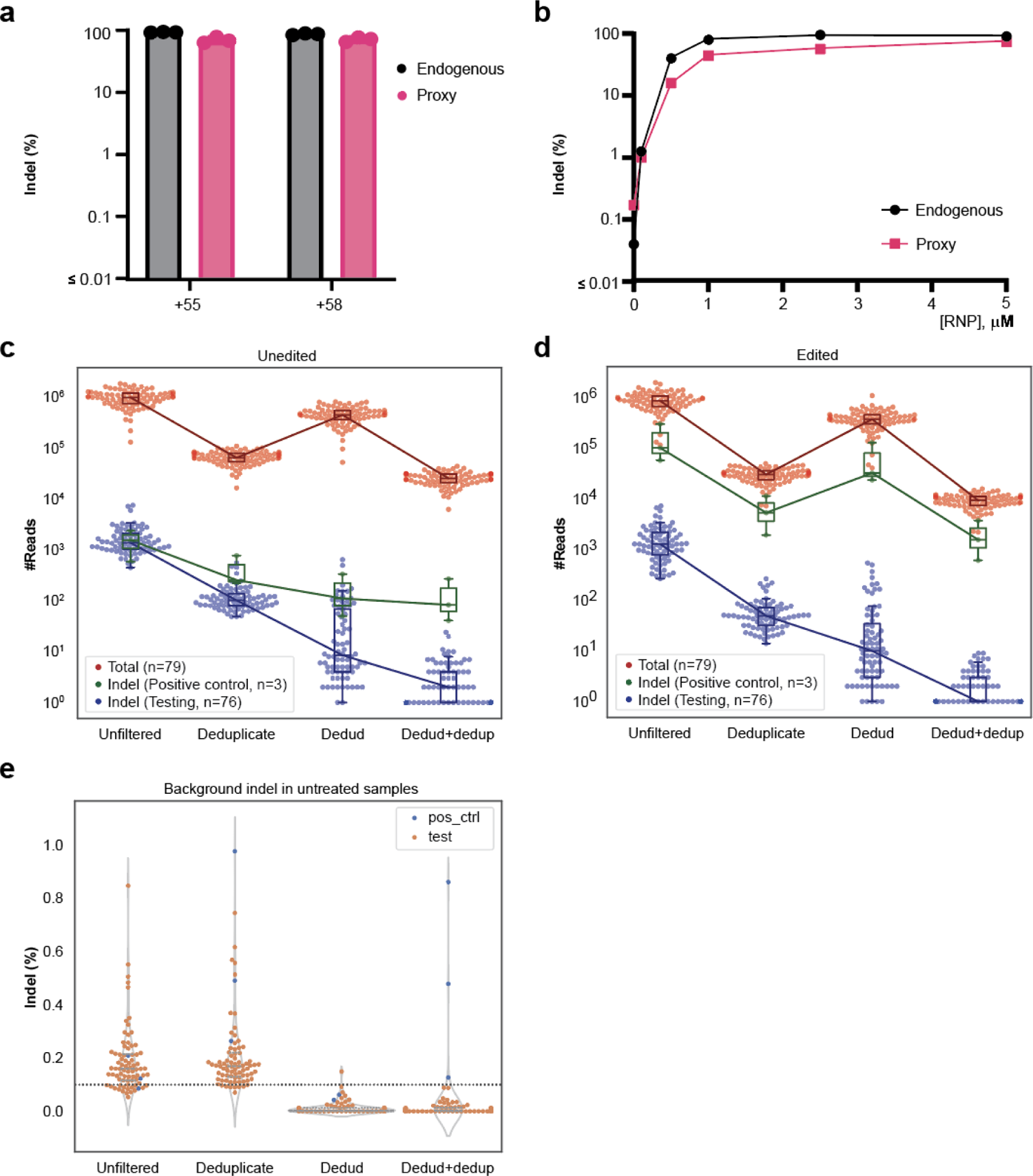
ABSOLVE-seq accurately and sensitively captures editing at integrant sequences. (A) Comparison of editing observed at endogenous vs. integrant loci at the on-target sequences for sgRNAs 1450 and 1617. (B) Evaluation of sensitivity in capturing true editing across indel frequencies. (C) Comparison of reads passing different filtering strategies in unedited and (D) edited samples. (E) Examination of background indel frequency in unedited samples with different filtering strategies. Dotted line marks the 0.1% indel threshold.

### ABSOLVE-seq enables high-throughput evaluation of candidate variant-associated off-targets

*In silico* off-target nomination using CRISPRme for sgRNA sg1617 previously identified the rs114518452 alternative allele (rs114518452-C) associated off-target site, which we then experimentally verified by performing Cas9 RNP editing of HSPCs from a donor heterozygous for rs114518452-G/C^10^. CRISPRme analysis of sgRNAs sg1617 and sg1450 against hg38 plus 1000 Genomes variants, with a cutting frequency determination (CFD) score^29^ ≥0.4, identified an additional 73 potential off-target sequences associated with genetic variants (**Supplementary Figure 3**). In total, we included in the ABSOLVE-seq integrant off-target lentiviral vector library 79 candidate sequences (**Table 1**): 2 on-target sequences, 1 variant-associated off-target found and verified in a previous analysis (variant rs114518452 associated, aka integrant 2-210530658_30_C23G), and 76 candidate off-target sequences (73 associated with genetic variant alternative alleles). Therefore, the 2 on-target sequences and integrant 2-210530658_30_C23G serve as internal positive controls for the experiment. By sequencing the plasmid pool, we observed that 97.5% of the reads contained one of the 79 expected candidate-barcode-candidate sequences. Each candidate had an average of 4.3e6 reads and 1.4e6 UMIs (**Supplementary Figure 4**).

**Table 1.**
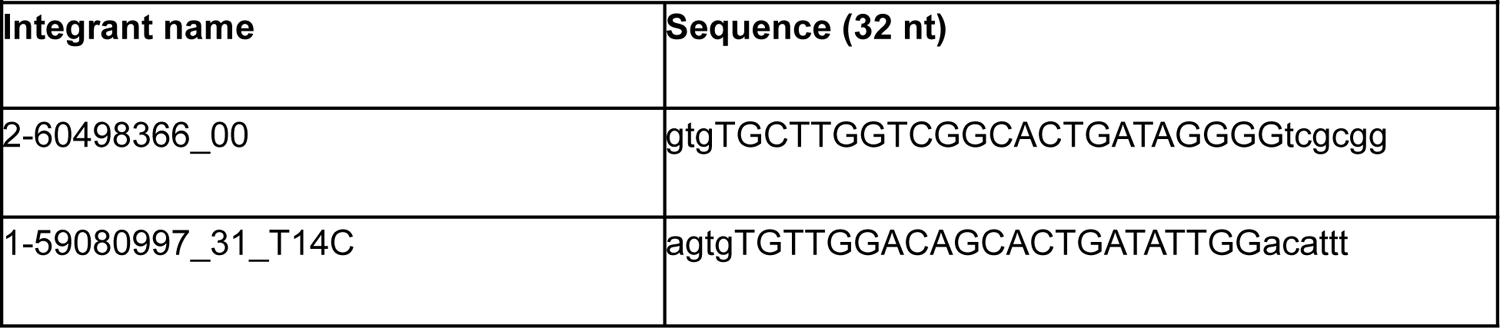
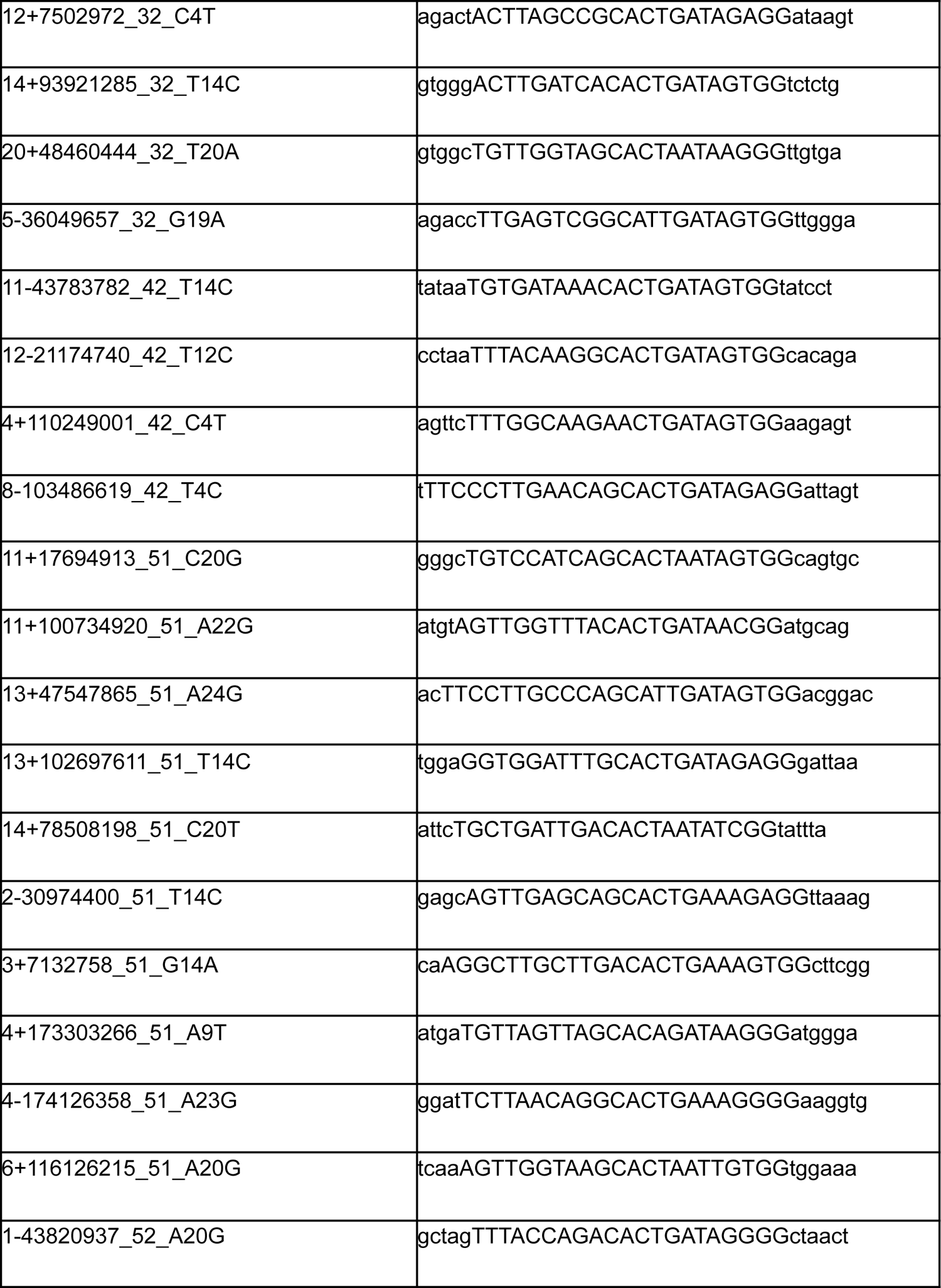
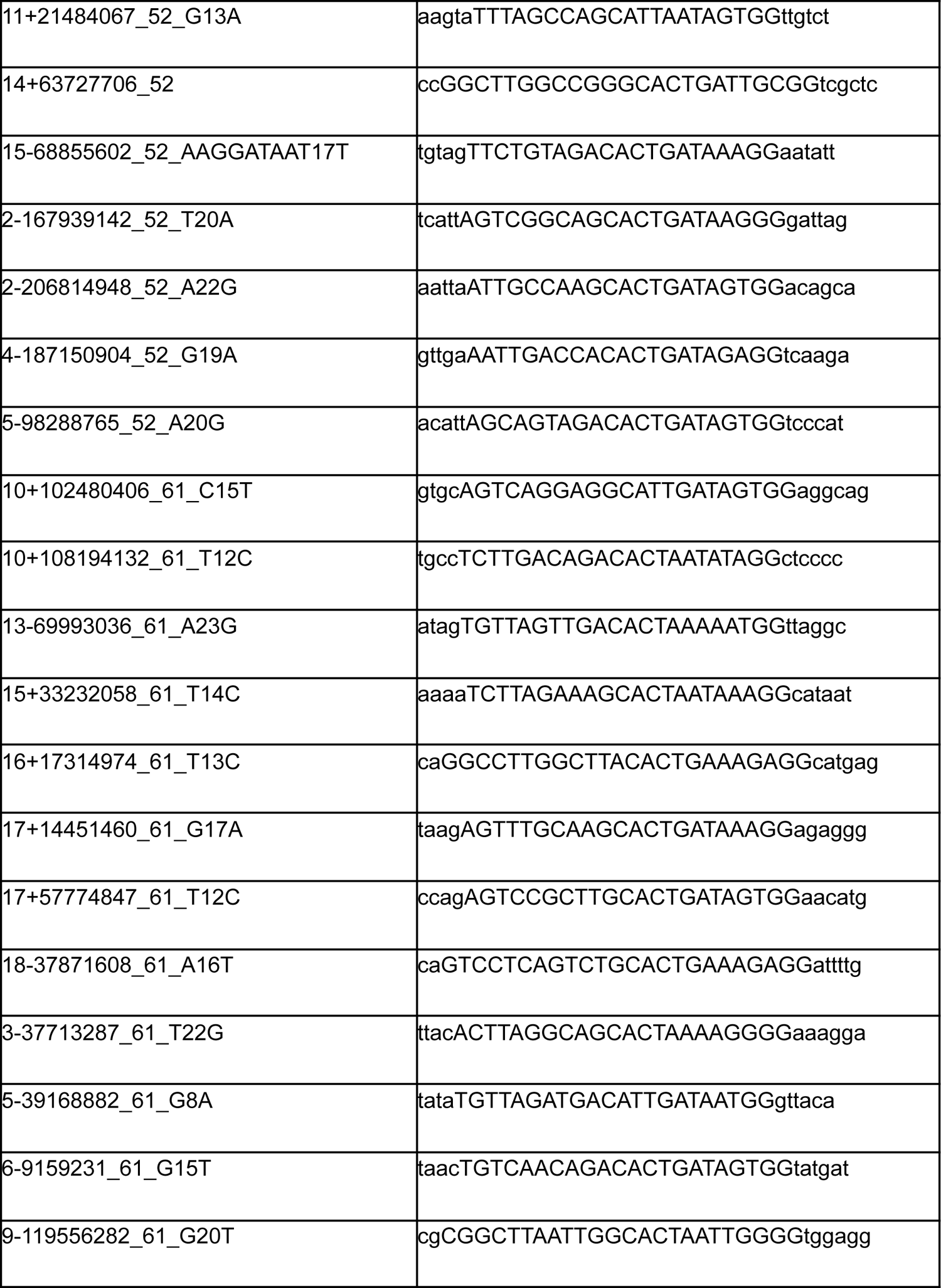
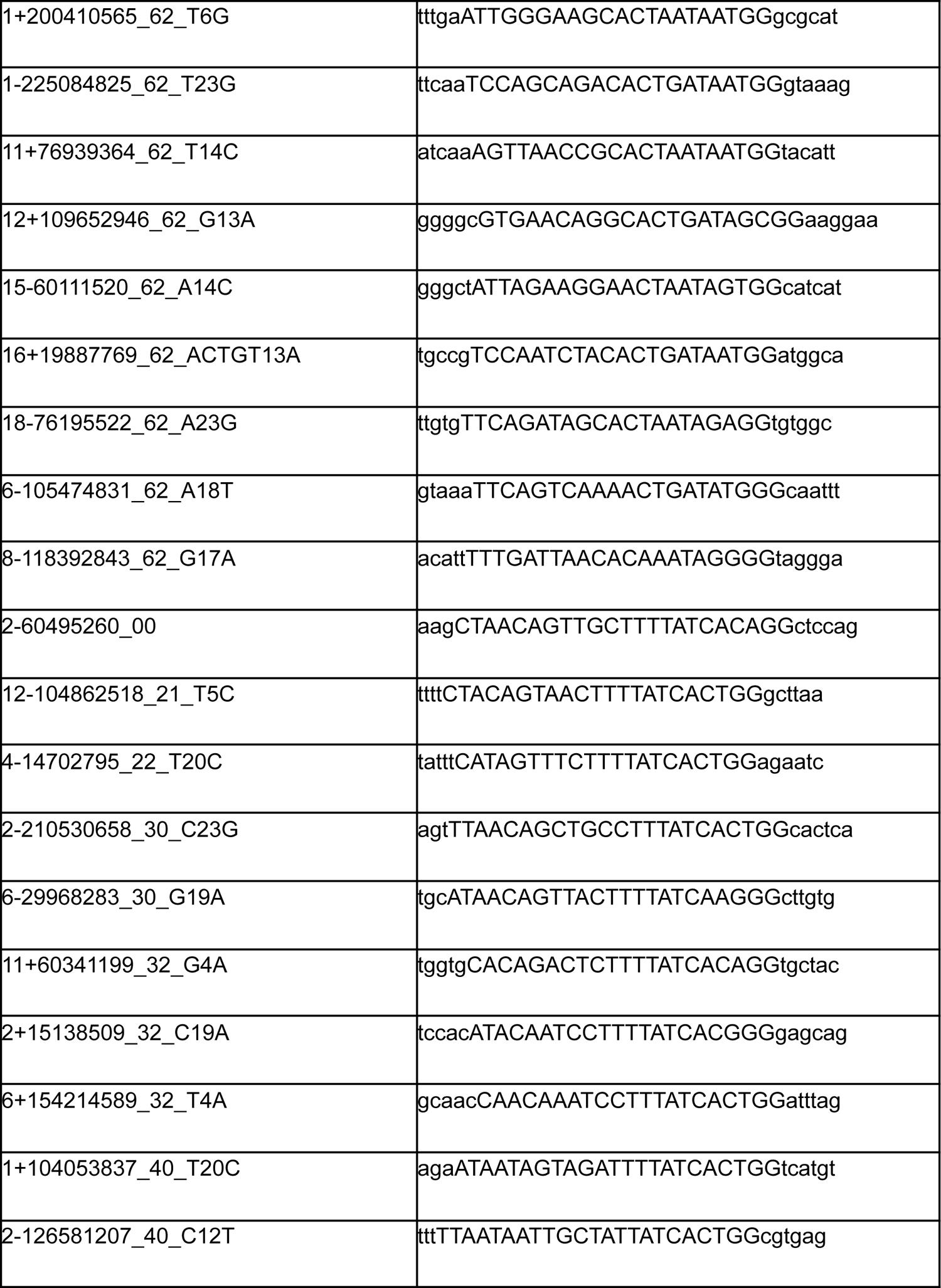
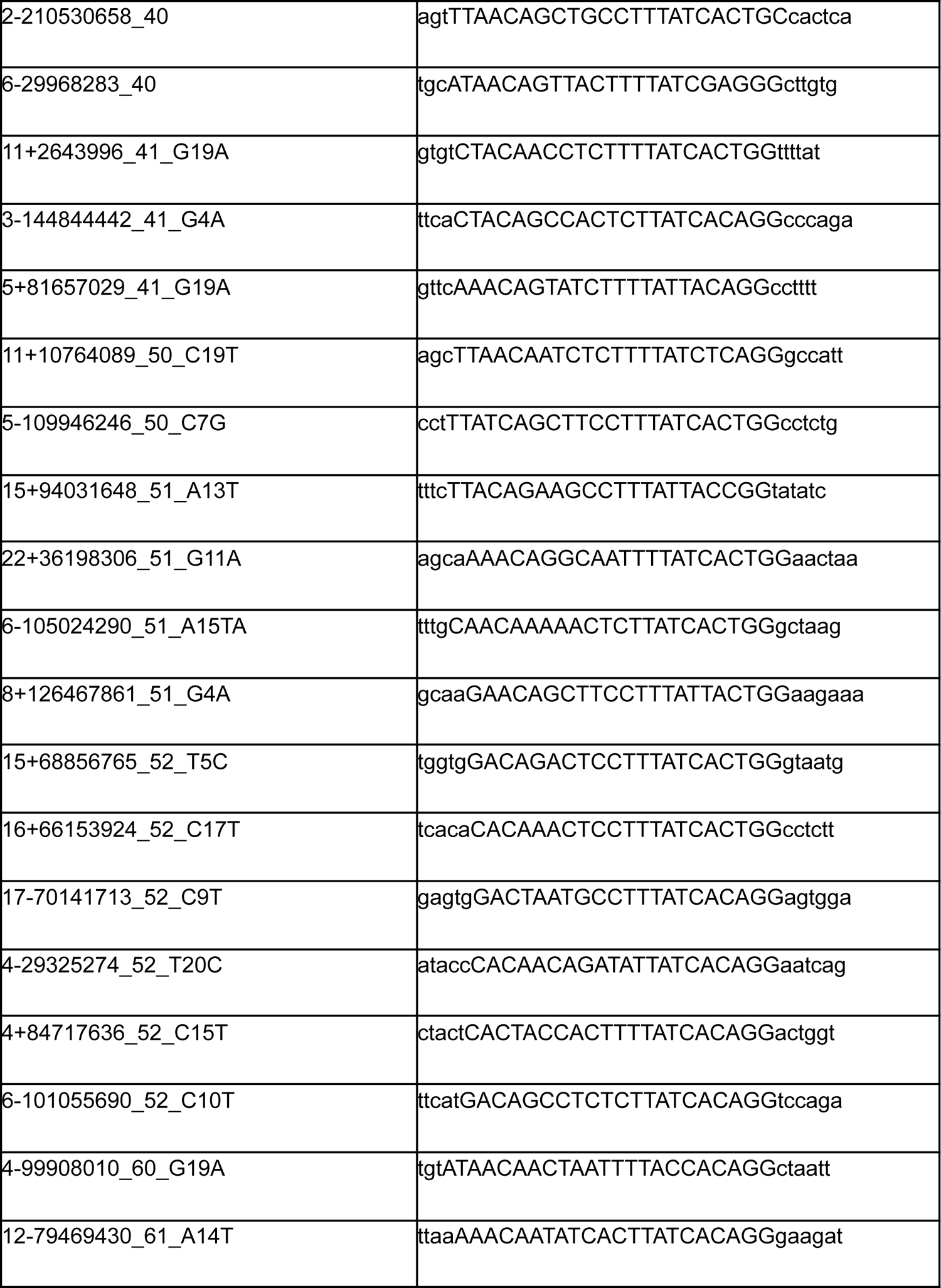
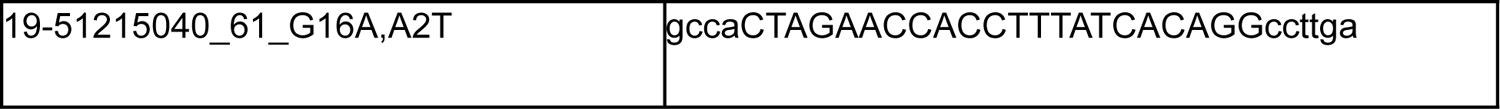
Integrant sequences included in the pooled library. These 32 nt integrants are based on genomic sequences from 5’ to 3’ flanking upstream (3 nt), candidate protospacer (20 nt), protospacer adjacent motif (3 nt), and flanking downstream (6 nt). Integrants are named as <chromosome> <strand><start position<_<Reference allele><position of variant within protospacer><alternative allele>_<number of mismatches compared to on-target><number of bulges when aligned to on-target>.

Following the experimental protocol described above, CD34+ HSPCs were transduced with the ABSOLVE-seq lentiviral library and either electroporated with RNP (edited) or not(unedited control). We observed mean transduction efficiency of 93.1% and vector copy number of 5.2 per diploid genome (**Supplementary Figure 2**). We isolated genomic DNA, amplified the ABSOLVE-seq integrant libraries, and sequenced 2-3 technical replicates per sample. As shown in **Figures 2C-D** (*Unfiltered* group), we obtained an average of 1.1e6 reads and 4.9e5 UMIs for each candidate off-target sequence for edited and unedited samples (**Supplementary Figure 4**). The reads from each sequencing run were demultiplexed using the candidate sequence barcode and aggregated into FASTQ files. These FASTQ files were then analyzed using CRISPResso2 for determining read coverage and indel frequency for each candidate sequence in our samples. We consider as indels all alleles composed of insertions and deletions overlapping a 1 bp window surrounding the expected SpCas9 cleavage site, as per CRISPResso default.

### Combining plasmid pool and integrant sequencing data reduces background noise

In our initial investigation, we focused on assessing background noise by calculating the ratio between reads carrying indels (*IndelReads*) and total reads (*TotalReads*). In unedited cells (**Figure 2E**, *Unfiltered* group), indel frequencies ranged from 0.05% to 0.85%, with an median editing frequency of 0.16%. These results are similar to published results for lentiviral integrants, such as Surro-seq, which reports an average indel percentage of 0.30% (ranging from 0% to 3.95%) in unedited samples^20^. Our observed background indel percentage, although comparable to established methods, surpasses the thresholds commonly used in the literature to distinguish relevant off-target editing events that would require risk assessment as part of preclinical assessment of a gene editing therapeutic candidate^25,30,31^. To explain this phenomenon, we hypothesized that these background indels might result from errors introduced during oligonucleotide synthesis, cloning of oligos to plasmid, sequencing library preparation, sequencing, and/or DNA carryover/cross-contamination among samples. These errors, when subsequently sequenced, can spuriously inflate the number of IndelReads.

To better understand the nature of these errors, we aligned reads from the plasmid pool to the expected construct. We found that 6.5% of reads did not match the expected construct sequence. Notably, 20% of these mismatched reads contained insertions or deletions at the junction of the oligo and plasmid vector backbone, suggesting errors during Gibson assembly, potentially due to aberrant resection and rejoining without complete homology. Additionally, we observed that deletions not overlapping the oligo/vector junction were predominantly 1 bp (69%), followed by 2 bp (12%) and 3 bp (8%), distributed throughout the read, suggesting expected oligo synthesis errors^32^ (**Supplementary Figure 5**). We also detected a constant and uniformly distributed error background of individual base pair mismatches compatible with Illumina sequencing errors^33^. Furthermore, 66% of the UMIs found in the plasmid pool are associated with a single barcode, and the degree of collision (defined as >1 UMI per barcode) ranges from 2 to 86 barcodes. We also observed some homopolymer UMI sequences and UMIs sequences with low complexity (**Supplementary Figure 6**).

To reduce background noise introduced by these errors, we explored four pre-processing and filtering strategies for read data to generate candidate-specific allele tables displaying indel and total read frequency. The first strategy, termed *Unfiltered*, utilizes all reads matching an expected barcode without any additional filtering. The second approach, *Dedup* (Deduplication), removes duplicate reads from the same parent DNA molecule and suppresses low-frequency errors by keeping only the most frequent read for each UMI. The third strategy, *Dedud*, retains only UMI sequences associated with at least one barcode-target sequence perfectly matching the design (**Supplementary Figure 7**). Furthermore, these high-fidelity reads are defined as those associated with a maximum of 2 barcodes and with UMI complexity (i.e., nucleotide entropy) above the 5th percentile. The fourth strategy, *Dedud+Dedup*, includes both filters, i.e. deduplication of high-fidelity reads. It is important to highlight that the *Unfiltered* and *Dedup* filtering strategies do not require sequencing data from the plasmid pool.

These filtering strategies progressively reduce the number of reads used for estimating indel frequencies, as shown in **Figures 2C** and **2D** for unedited and edited samples, respectively. Since *Dedup* relies on the number of UMIs for indel quantification, the total amount of data points used for inference is strongly reduced. On average, only 5% of the *Unfiltered* reads are retained (Avg. *TotalReads Dedup*: 96,931 reads). A similar data loss is observed in unedited samples for noisy reads labeled as indels (Avg. *IndelReads Unfiltered*: 1,769 reads, Avg. *IndelReads Dedup*: 138 reads), resulting in comparable median background indel rates of 0.16% for *Unfiltered* and 0.17% for *Dedup* filtering among 79 candidates (**Figure 2E**). The *Dedud* strategy retains on average approximately 44.8% of the total (*Unfiltered*) reads (Avg. *TotalReads Dedud*: 855,667 reads), but only on average 1.9% of the indel reads (Avg. *IndelReads Dedud*: 2,501 reads), lowering the average background noise to 0.002%. When deduplication is combined with the *Dedud* strategy, on average 1.8% (Avg. *TotalReads Dedup+Dedud*: 35,039 reads) of the total reads and 0.01% (Avg. *IndelReads Dedup+Dedud*: 84 reads) of the indel-carrying reads pass the filtering, leading to an estimated indel background of 0.004% (**Figure 2E**). As shown in **Figures 2C** and **2D**, the different filtering strategies induce similar *TotalReads* loss in the edited samples as in the unedited samples. As expected, within *IndelReads* across all strategies considered, the positive control integrants have a higher number of *IndelReads* compared to the 76 unverified candidates.

### Estimating Editing Activity and Enhancing Statistical Power through Dedud Filtering

The frequency tables serve as the response variable for estimating indel proportion. Importantly, this model allows us to weigh samples according to their coverage, providing a more accurate estimation. This estimation is performed using a generalized linear model for binomial responses, which is estimated using a maximum penalized likelihood approach^34,35^. We use samples’ annotation data, such as donor, replicate, and editing status (edited/unedited), as covariates. This enables us to identify potential confounders, such as donor-specific effects. Editing at a candidate sequence is estimated as the marginal Δindels, which is the difference between indels in the edited and unedited sample. This allows us to intrinsically consider regions with varying intensities of background noise. Δ%indels estimation corresponds to the transformation of the coefficient for the Editing variable in the regression model. For this, we can use asymptotic results for calculating confidence intervals and testing enrichment significance using the Wald test^36^.

The filtering strategies applied during data preprocessing significantly impact the number of reads retained for inference, which, in turn, affects the reliability and detection rate of ABSOLVE-seq. To explore this relationship, we conducted a Monte Carlo-based power analysis. Specifically, we focused on data from unedited samples and calculated donor-specific background %indel (denoted as p*_ij_*, where *i* corresponds to candidate and *j* to donor). Next, we simulated *IndelReads* for the edited samples using a binomial distribution (N,p), adjusting the success probability to p*_ij_* + (0.1% or 0.01%) and setting the number of trials N to experimental *TotalReads* of the edited samples. Merging the simulated data with experimental data for the unedited samples, we estimated Δ%indels and tested its significance. These in-silico experiments were repeated 1000 times for each sequence and filtering strategy, recording the percentage of experiments resulting in statistically significant Δ%indels estimates.

In **Figure 3A**, we compare editing frequency estimates for three positive control integrant sequences using the *Unfiltered* and *Dedud* filtering strategies: sgRNA1450 on-target (2-60498366_00), sgRNA1617 on-target (2-60495260_00), and verified off-target 2-210530658_30_C23G. The marginal increment in %indels in edited samples was significant for all three positive controls with both strategies, and *Unfiltered* and *Dedud* produced similar Δ%indels estimates (2-60498366_00: 76.88% for *Unfiltered* and 79.12% for *Dedud*; 2-60495260_00: 80.59% for *Unfiltered* and 83.28% for *Dedud*; 2-210530658_30_C23G: 4.29% for *Unfiltered* and 3.89% for *Dedud*). Background %indels in unedited samples were consistently lower with *Dedud* vs. *Unfiltered* (2-60498366_00: 0.04% vs. 0.09%; 2-60495260_00: 0.03% vs. 0.13%; 2-210530658_30_C23G: 0.008% vs. 0.1%). For 2-210530658_30_C23G, donor-specific estimates showed high variability. Overall, *Dedud* reduced background indels while maintaining similar editing frequencies for positive controls.

**Figure 3.**
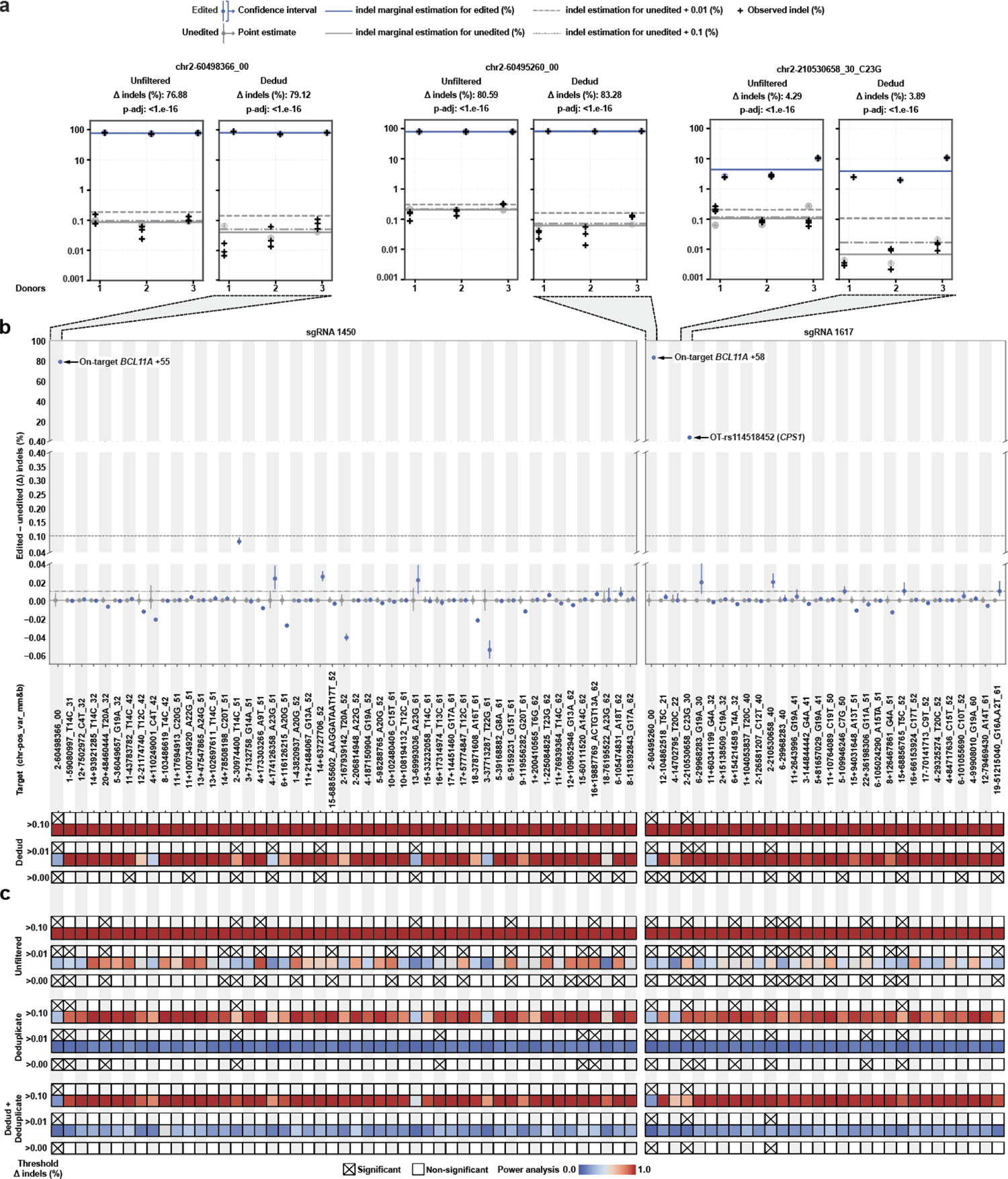
Computational filtering and statistical modeling of sequencing data enables identification of off-targets with high sensitivity and power. (A) Editing estimates for the positive control sequences using the *Unfiltered* and *Dedud* filtering strategies. (B) Editing estimates for all candidate sequences in the lentiviral pool using the *Dedud* filtering strategy. Candidates are ordered by increasing number of mismatches followed by bulges to the two on-target sequences. (C) Comparison of significant editing calls at various thresholds via different filtering strategies with associated power.

Figure 3B presents the marginal Δ%indels estimates alongside their corresponding confidence intervals for all integrant sequences using the *Dedud* filtering strategy. Notably, only the three positive controls highlighted in Figure 3A exhibit a significant Δ%indels > 0.1%. Eight additional integrant sequences have estimates for Δ%indels > 0.01% and ≤ 0.1%, while nine additional ones have statistically significant estimates with Δ%indels > 0.0 and ≤ 0.01%.

In Figure 3C, we compare ABSOLVE-seq performance among filtering strategies and evaluate significant off-target calls using three increasingly stringent criteria: statistical significance (FDR adjusted p-value for Δ%indels < 0.05), statistical significance combined with Δ%indels > 0.01% and ≤0.1%, and statistical significance combined with Δ%indels > 0.1%. Notably, all filtering strategies successfully identify the three positive controls as significant. However, *Dedud+Dedup* detect one additional significant off-target sequence.

Using the Dedup strategy, we identified 19 significant sequences, each with a Δ%indels > 0.01%. Among these, 8 sequences, which include 3 positive controls and 5 candidates, demonstrated an estimated Δ%indels > 0.1%. Among all filtering strategies, the Unfiltered approach yielded the highest number of significant sequences, totaling 33. Of these, 32 sequences exhibited a Δ%indels > 0.01%, while the three positive controls and 12 candidate sequences showed a Δ%indels > 0.1%.

As depicted in Figure 3B**/C**, both the *Unfiltered* and *Dedud* strategies consistently provide indel frequency data that ensures a statistical power greater than 0.99 for Δ%indels ≥ 0.1. However, the *Dedup* and *Dedup+Dedud* criterias encounter challenges. Multiple candidate sequences within their final datasets fail to achieve a satisfactory level of statistical power (>80%). Remarkably, the *Dedud* processed frequency table allows for the detection of Δ%indels ≥ 0.01 in 69 out of the 79 candidate sequences. Two out of the ten sequences that do not consistently have sufficient statistical power for detecting Δ%indels ≥ 0.1 are the on-target integrants, for which the Δ%indels in the edited sample are much larger. We surmise this is due to having inflated background %indels in unedited samples for these integrant sequences, caused by carryover and cross-contamination with the respective highly edited samples during library prep or sequencing steps^37,38^.

### Insights and guidelines for maximizing ABSOLVE-seq precision and statistical power

ABSOLVE-seq’s optimal filtering strategy, *Dedud*, relies on sequencing data from both the plasmid pool and unedited and edited cells to selectively detect and filter out low-quality reads, thereby reducing background noise and improving statistical power. To better understand how the quality of the plasmid pool and sequencing depth impact the precision and sensitivity of ABSOLVE-seq, we developed a comprehensive statistical model to predict UMI yield as a function of sequencing depth. Using UMI recapture frequency (**Supplmentary Figure 8**) from three sequencing runs and capture-recapture models (Chao.1 estimator), we estimated 36,851,341 unique unfiltered UMIs in the plasmid pool, with 24,790,143 (67.3%) surviving *Dedud* filtering. We integrated these values with the cumulative number of unique UMIs detected in the plasmid sequencing runs to estimate a nonlinear asymptotic regression model.

Our experimental sequencing depth analysis and model results predict that approximately 40% of the UMIs in the plasmid pool have been observed, with 42% detected in *Dedud*-filtered data and 36% in Unfiltered data (Figure 4A). It is important to note that the estimated richness of the plasmid pool by the Chao.1 estimator represents a lower bound, suggesting that the true proportion of observed UMIs might be lower. Furthermore, our model indicates that around 1.5 billion reads from the plasmid pool would be necessary to recover 90% of the total UMI population.

**Figure 4:**
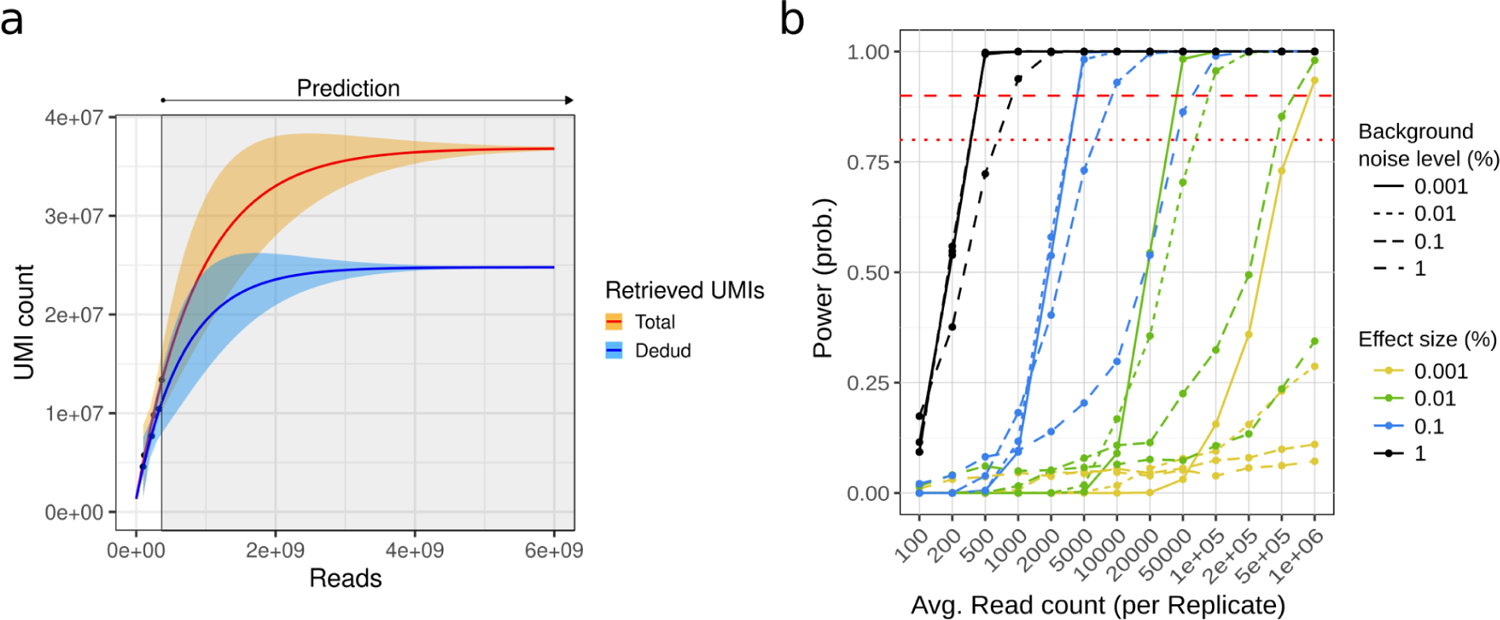
Modeling of sequencing depth, UMI recovery, and statistical power provide guidelines for ABSOLVE-seq experimental design optimization **(A)** Non linear model for the relationship between sequencing depth of the plasmid pool and the number of recovered UMIs. The orange region represents the total retrieved UMIs, while the blue region shows the UMIs employed by Dedud filtering. The left part of the plot shows experimental data, while the right (Prediction) is the predicted number of UMI for increasing sequencing depths. **(B)** Results of the simulation study investigating the impact of indel background noise and target effect size on statistical power across varying sequencing depths. Each line represents a different combination of background noise levels (solid: 0.001%, fine dashed: 0.01%, dashed: 0.1%, long dashed: 1%) and target effect sizes (yellow: 0.001%, green: 0.01%, blue: 0.1%, black: 1%). Each dot represents the statistical power calculated on 1000 simulation experiments.

To provide guidelines and best practices for future users, we conducted a simulation study to determine the data requirements—either in terms of reads or UMIs, depending on the filtering strategy—needed to detect specific target levels of Δ%indels with high probability under varying background noise scenarios. Using the generative model developed for statistical power calculation, we created in-silico unedited and edited samples with four distinct levels of background noise and effect sizes (1e-3, 1e-2, 1e-1, 1e0). For each setting, we considered 13 sequencing depths ranging from 1e2 to 1e6 and adopted a simplified experimental design consisting of a single donor and three technical replicates. We performed 1,000 simulations for each combination of settings, calculating the statistical power by assessing how often our model correctly identified a statistically significant positive Δ%indel.

Our results shown in Figure 4B highlight the specific sequencing depths required to achieve adequate power, depending on noise levels and target effect sizes, with both factors having a similar magnitude of impact. For background noise levels estimated using the *Unfiltered* strategy (0.1%) and *Dedud* strategy (0.01%), the sequencing depths needed for reliably detecting a Δ%indel > 0.1% were relatively similar: 5,000-10,000 reads for *Unfiltered* and 2,000-5,000 reads for *Dedud*. However, detecting smaller Δ%indel sizes, such as 0.01%, required significantly higher sequencing depths—200,000-500,000 reads with noise at 0.1% (*Unfiltered*) and 50,000-100,000 reads with noise at 0.01% (*Dedud*).

This deeper characterization of ABSOLVE-seq performance for varying level of background noise and target effect sizes provides future users with the ability to tailor their sequencing and experimental design to match their precision and statistical power goals effectively. Our findings suggest that when sufficient data are available, ABSOLVE-seq can reliably identify off-target editing events rarer than 0.1%. This capability is crucial for loci of particular biological and clinical relevance, such as cancer-associated variants, where even low-frequency off-target editing can have significant implications.

## DISCUSSION

ABSOLVE-seq addresses a critical limitation in experimental verification of candidate off-target sites that may compromise the specificity of therapeutic gene editing. It enables scalable testing of genetic variant associated candidate off-target sequences in genomic context with relevant editor delivery mechanisms and cellular environments when primary cells of all relevant genotypes are not available for direct verification (Figure 1). ABSOLVE-seq facilitates examination of the specificity of gene editing reagents during preclinical therapeutic development by artificially constructing in primary cells gRNA-dependent genetic landscapes that present elevated risks for off-target effects, thus mimicking the risk present in a larger population of individuals with varying genotypes.

A key advantage of ABSOLVE-seq lies in its ability to multiplex the assessment of numerous potential off-target sequences within a single experiment including internal controls. To demonstrate this throughput, we constructed a lentiviral library harboring a pool of 79 candidate off-target sequences for sg1617 and sg1450, two clinically relevant gRNAs targeting Cas9 to edit the +58 and +55 *BCL11A* erythroid enhancer for β-hemoglobinopathies. CD34+ HSPCs from 3 donors were transduced with the ABSOLVE-seq lentiviral library and edited using a clinically relevant Cas9 RNP complex and electroporation protocol. We demonstrate that the ABSOLVE-seq assay closely reflects editing activity observed at endogenous sites across a wide range of indel frequencies (Figure 2B). This finding highlights the assay’s reliability to capture true editing events.

Our regression-based estimation procedure directly estimates indel frequency in edited samples while accounting for sequence-specific background noise and experimental design factors such as donor variability. By utilizing *Unfiltered* indel frequency and coverage data, we demonstrate that our integrant sequences serve as accurate predictors at endogenous sites (Figure 3A). Statistical power analysis on the *Unfiltered* integrant datasets indicates that the sequencing depth and data quality are sufficient to detect biologically relevant %Indel increments of 0.1% or more with high confidence. However, we observed up to 1% indel frequency in unedited samples, which negatively impacts the precision and fidelity of the assay.

We developed four filtering strategies to reduce background noise, allowing users to balance sequencing depth and assay sensitivity. These strategies combine a quality-based filter (*Dedud*) with a deduplication (*Dedup*) step. *Dedud* leverages high-fidelity reads matching the expected structure design and carrying a UMI sequence found in the plasmid pool to be associated with up to two integrant sequences. *Dedud* specifically reduces the number of *IndelReads* in the unedited samples while preserving coverage of integrant sequence reads. On the contrary, in unedited samples, *Dedup* depletes the same proportion of indel and total reads from the unfiltered datasets, showing a lack of selectivity for false *IndelReads* (Figure 2C**,D**). Additionally, since *Dedup* quantification is based on the number of UMIs, it relies on significantly fewer data points compared to *Unfiltered* and *Dedud*, which are based on read counts. As a result, *Dedup* experiences a substantial reduction in integrant coverage, with an average data amount that is 9.7e4 (UMIs), 19, and 8 times smaller than that of *Unfiltered* (1.9e6 reads) and *Dedud* strategies (8.6e5 reads), respectively. As demonstrated in Figure 2E, *Dedud* significantly reduces the level of background indels (0.002%) compared to *Unfiltered* data (0.16%) and *Dedup* alone (0.17%), while combining *Dedup*+*Dedud* increases noise to 0.004%.

Regardless of the filtering method used (*Unfiltered* or *Dedud*), positive control estimates closely match those from independent experiments, supporting the assay’s accuracy (Figures 2**,3A**). Notably, the *Dedud* strategy significantly lowers background noise not only on average across all candidate sequences but also consistently in each individual sample analyzed. This consistent reduction in noise levels demonstrates the robustness of the *Dedud* filtering approach, providing reliable performance enhancements for every analyzed sample when compared to the *Unfiltered* strategy.

Our modeling approach allows us to estimate donors’ marginal distributions, an essential tool to highlight significant donor variability, as we observed for integrant 2-210530658_30_C23G, for which donors exhibit editing estimates ranging between 3% and 10%. In all settings, the two on-target integrants reported higher levels of background noise, attributable to potential low-level DNA carry-over/cross contamination between samples with extremely high levels of editing and their unedited counterparts^40^.

The improved signal-to-noise ratio achieved by *Dedud* translates to a significant boost in assay sensitivity. This is evident in Figure 3B, where the assay exhibits high statistical power (>80%) to detect editing effects with a Δ%indels threshold of 0.01% for the majority of the candidate sequences (68 out 79). This threshold is tenfold lower than the current standard in the field, allowing ABSOLVE-seq to identify previously undetectable low-level off-target editing events. Among the 20 candidate sequences with statistically significant positive Δ%indels identified using *Dedud*, only the positive controls exceeded the 0.1% threshold. Eight additional sequences exhibited editing effects between 0.1% and 0.01%, highlighting the assay’s ability to detect low-level off-target activity, although the biological and clinical importance of these sequences may be uncertain.

ABSOLVE-seq, with the *Unfiltered* strategy, identifies a higher number of potentially edited sequences. This includes the three positive controls and an additional 12 candidates exceeding a Δ%indels threshold of 0.1%. While it also detects 17 more sequences with Δ%indels exceeding 0.01%, unlike the *Dedud* strategy, the *Unfiltered* approach suffers from limitations in elevated background noise. These shortcomings reduce the assay’s statistical power to detect subtle editing effects (Δ%indels > 0.01%) reliably. Consequently, some of the additional sequences identified by the *Unfiltered* strategy appear likely to be false positives.

The accuracy and sensitivity of ABSOLVE-seq are contingent on both the experimental background noise and the sequencing depth of the plasmid pool, as well as unedited and edited samples. As illustrated in Figure 4, we developed a statistical model that estimates the richness of the plasmid pool and the UMI yield as a function of sequencing reads by analyzing UMI capture-recapture patterns. With defined target effect sizes, our model allows us to devise experimental plans and determine the optimal sequencing depths needed for ABSOLVE-seq to meet requirements while avoiding excessive sequencing. The current threshold of 0.1% Δ%indels is often used to identify true positive editing, but this benchmark is based more on the technical limitations of existing sequencing technologies than on biological rationale. We foresee the future development of ABSOLVE-seq incorporating different thresholds for biological significance, tailored to the potential impact of off-target editing. More stringent criteria might be applied to candidate off-target sites within critical regulatory regions of cancer-associated genes, for instance. Importantly, any conclusion about the absence of significant off-target editing should always be supported by a power analysis that confirms the experimental data is sufficient to detect a target effect size if present. While ABSOLVE-seq enables scalable verification of candidate off-target sequences in relevant cellular and gene editor delivery contexts, there remain some limitations to the method.

We observed higher background indels in unedited samples for the on-target sequences as compared to candidate off-targets. We hypothesize that the higher background of the on-target integrant sequences is due to low-level contamination of edited reads in unedited samples. Of note, these background indel frequencies remain ∼3 orders of magnitude lower than indels observed in edited samples. Practically, this seemingly heightened background does not result in risk of false positive assignment since for these integrants the true signal is readily apparent with orders of magnitude higher editing in the edited samples. In contrast, for the 76 previously unverified candidates where we observed indels in the edited samples <1% for each candidate, the background indels in unedited samples were 0.01% (median, with range 0.001 to 0.02%), demonstrating the sensitivity of the method.

Here we utilized an analytical filtering process to remove errors from the library to improve power to detect off-target editing. Improved library synthesis, cloning, library preparation and sequencing methods to improve the fidelity of sequences comprising the plasmid pool could reduce the need for this analytical filtering step which comes at a modest cost of reduced read utilization.

Another limitation is that the ABSOLVE-seq integrants do not precisely mimic the chromatin context of the candidate sequences in relevant cell types. It is well known that lentiviral integrations tend to be concentrated in open chromatin regions^41,42^. Therefore the assay would be expected to overestimate the editing frequency of candidate off-targets that may naturally reside in closed chromatin regions. Given a main use case for the assay could be demonstrating that certain candidate variant-associated off-target sites do not demonstrate significant editing with adequate power to detect effects at a given biologically relevant threshold, this tendency of the assay to overestimate editing may be acceptable to users.

For this analysis, we utilized nomination of candidate off-target sites based on mismatch and bulge thresholds combined with CFD score, a simple to calculate score that has been shown to identify experimentally verified off-target sequences. Still we found that 76 of 76 candidate off-target sites failed to be verified at a threshold of 0.1% Δindels. As models of off-target editing, guided by free energy and deep learning models, continue to improve, we might include smaller sets of candidate off-targets with higher pre-test probability to be verified.

Low-level off-targets remain a challenge in the field of gene editing specificity analysis, with the biological significance uncertain. Integrating ABSOLVE-seq (or other off-target assays) with functional assays could provide a more in-depth understanding of off-target editing consequences, extending beyond mere detection to understanding biological safety.

Future efforts to enhance ABSOLVE-seq’s modeling framework by considering allele frequency tables as a response variable instead of indel counts might take advantage of more subtle editing signals that span larger regions or involve complex editing patterns. Modeling allele frequencies could improve accuracy and sensitivity by capturing these intricate changes, while also providing valuable insights into the molecular pathways involved in DNA repair post-editing. This approach has the potential to deepen our understanding of how different editing outcomes are influenced by the cellular context and improve the precision of future off-target prediction tools.

Notably, ABSOLVE-seq can be adapted to evaluate non-nuclease editing tools like base editors and prime editors^43^. These tools hold great therapeutic potential, but their unique off-target profiles necessitate careful characterization. ABSOLVE-seq can be a valuable asset in this context, paving the way for a standardized platform to assess off-target editing across diverse editing modalities. By addressing these aspects, ABSOLVE-seq can become an indispensable tool for ensuring the safety and efficacy of next-generation gene editing therapies and fill an important current gap in providing a scalable robust assay for experimental verification of candidate off-targets associated with genetic variants in a relevant cellular context.

## METHODS

### Plasmid design and vector production

The LV vector was generated from 3^rd^ generation pLV-SFFV-eGFP-WPRE by first inserting the synthesized double-stranded oligonucleotide GCGATCGCTTAATTAACTCTTTCCCTACACGAC GCTCTTCCGATCTCACCGGAGACGGTTCTCTGGATGATTGTACGTCTCTTCGAAGATCGGAAGAGCACACGTCTGAACTCCAGTTAATTAAGCGATCGC with added compatible single-stranded overhangs into the XhoI cut site immediately upstream of the SFFV promoter. This insert contains RE sites for PacI and AsiSI on both ends and a central BsmBI cut site for subsequent cloning steps. The insert for the second cloning step was generated by annealing of Forward 5’-CTACACGACGCTCTTCCGATCTNNNNNNNNNNNNNNNNCCAACCTCATAGAACACTCATCC-3’ and Reverse 5’-AGACGTGTGCTCTTCCGATCT-protospacer+PAM[32](complement)-XXXXXXXXX-GGATGAGTGTTCTATGAGGTTGG-3’ followed by PCR amplification with Primers For-5’-CTCTTTCCCTACACGACGCTCTTCCGATCT-3’ and Rev 5’-AGACGTGTGCTCTTCCG-3’.

N16 indicates a randomized UMI sequence, X9 a defined barcode specific to the candidate sequence. The candidate sequence consists of the protospacer and protospacer adjacent motif (PAM) sequence (23 nt) plus genomic flanking sequence (3 nt upstream of protospacer and 6 nt downstream of PAM) (see **Table 1**). The barcode is positioned at the PAM-distal end of the protospacer+PAM sequence.

This insert was cloned into the BsmBI site of the digested LV-vector via Gibson assembly (New England Biolabs, cat# E5510) with 0.1 pmol of digested vector and 0.5 pmol of insert. Representation of the different off-target site sequences and barcode complexity was ascertained by Sanger sequencing of individual clones and amplicon sequencing the pool. Virus supernatants were produced by transient transfection of 293T cells using the LV-plasmid and packaging plasmids pGag-Pol, pRSV-REV, pCMV-VSVg and pAdVAntage (pAdVA). Viral supernatants were harvested after 48h and concentrated 500-fold via ultracentrifugation.

### Transduction and RNP electroporation

Cryopreserved human CD34+ HSPCs from mobilized peripheral blood of deidentified healthy donors were obtained from Fred Hutchinson Cancer Research Center, Seattle, Washington. CD34+ HSPCs were thawed and cultured into Stem Cell Growth Medium (SCGM), GMP grade (CellGenix, cat#, 20806-0500) supplemented with 100 ng ml-1 Preclinical Thrombopoietin (TPO) (CellGenix, cat# 1417-050), 100 ng ml-1 Preclinical Stem Cell Growth Factor (SCF) (CellGenix, cat# 1418-050) and 100 ng ml-1 Preclinical FMS-like Tyrosine Kinase 3 Ligand (FLT3L) (CellGenix, cat# 1415-050). 24-48 hours post-thaw, CD34+ HSPCs were transduced with 10 µL of virus, 200 μg/mL F108 (Sigma, cat# O7579), and 8 μM Cyclosporin H (Sigma, cat# SML1575) in SCGM supplemented with cytokines as described above at a concentration of 1 million cells/mL. 16-18 hours after transduction, HSPCs were electroporated using Lonza 4D nucleofector. 300 pmol (15 μM) of sg1617 was mixed with 100 pmol (5 μM) of 3xNLS-SpCas9-HiFi protein and added electroporation buffer (Lonza 4D, cat# V4XP-3032) up to 5 μl in one tube, 300 pmol (15 μM) of sg1450 was mixed with 100 pmol (5 μM) of 3xNLS-SpCas9-HiFi protein and added electroporation buffer up to 5 μl in another tube. After 15 min of incubation at room temperature, RNP from the two tubes was mixed. 100,000 HSPCs were prepared and resuspended in 10 μl of electroporation buffer, RNPs were mixed with cells and transferred to a cuvette (Lonza 4D, cat# V4XP-3032). Electroporation was performed using the program EO-100 with Lonza 4D nucleofector. After electroporation, cells were washed twice and cultured for an additional six days in an erythroid differentiation medium (EDM) consisting of IMDM (GibcoTM, 12440061) supplemented with 330 μg/ml holo-human transferrin (Sigma, T0665-1G), 10 μg/ml recombinant human insulin (Sigma, 19278-5ML), 2 IU/ml heparin (Sigma, 19278-5ML), 5% human solvent detergent pooled plasma AB (Rhode Island Blood Center), 3 IU/ml erythropoietin (AMGEN, 55513-144-10), 10^-6^ M hydrocortisone (Sigma-Aldrich, H0135), 100 ng/ml human SCF (CellGenix, 1418-050) and 5 ng/ml of recombinant human IL-3 (PEPROTECH, 200-03). Seven days after transduction, an aliquot of cells was resuspended in PBS and analyzed on a BD LSR II to measure transduction frequency through GFP expression. Data were analyzed using FlowJo^TM^ software. The remainder of the cells were used for genomic DNA isolation.

### Droplet digital PCR

Genomic DNA was isolated using the Blood and Tissue Kit (Qiagen cat# 69506) according to the vendor’s recommendations. 50 ng of genomic DNA was used for each assay. ddPCR was performed using Biorad QX200 AutoDG droplet digital PCR system. ddPCR supermix (no dUTP) (BioRad, cat# 1863023) and HindIII (NEB, cat# R3104S) were added in the reaction. Lentivirus vector copy number assay: Forward TACTGACGCTCTCGCACC, Reverse TCTCGACGCAGGACTCG, Probe ATCTCTCTCCTTCTAGCCTC. Albumin reference assay: Forward GCTGTCATCTCTTGTGGGCTGT, Reverse ACTCATGGGAGCTGCTGGTTC, Probe CCTGTCATGCCCACACAAATCTCTCC. After droplet generation, PCR followed the cycling conditions: 1 cycle of 95°C for 10 min, 40 cycles of 94°C for 30 sec, 60°C for 60 sec, 1 cycle of 98°C for 10 min, held at 4°C, ramp rate of 2°C per second. PCR products were read by the droplet reader and the results analyzed using the QuantaSoft software.

### Endogenous sequencing library preparation

The *BCL11A* enhancer +58 and +55 on-target sequences from the transduced and edited cells were amplified with KOD Hot Start DNA Polymerase (EMD-Millipore, 71086-31) and corresponding primers: +58 forward AGAGAGCCTTCCGAAAGAGG, +58 reverse GCCAGAAAAGAGATATGGCATC, +55 forward ACAGTGATAACCAGCAGGGC, +55 reverse GATGCAATGCTTGGAGGCTG. In a parallel experiment with a heterozygous donor carrying the alternative allele rs114518452-C, CD34+ cells were electroporated with 3xNLS-SpCas9-HiFi:sgRNA1617 and sg1450 as described above in three replicates. The endogenous OT-rs114518452 off-target sequence was amplified with KOD Hot Start DNA Polymerase with primers OT-rs114518452 forward TAAGATTCTTTTGGTTCTGGCT, OT-rs114518452 reverse AGAGAGGCAGTATTTACGATGC. The cycling conditions were 95 °C for 3 min; 30 cycles of 95 °C for 20 s, 60 °C for 10 s, and 70 °C for 10 s; 70 °C for 5 min. Locus-specific PCR products were indexed and sequenced with paired-end ∼150 bp reads on an Illumina HiSeq, NovaSeq, or MiniSeq instrument.

### Lentiviral integrant sequencing library preparation

The integrant cassette including UMI and barcode was amplified from genomic DNA with KOD Hot Start DNA Polymerase (EMD-Millipore, 71086-31) with two sets of PCR primers as below. Forward-outer: ACAGTGCAGGGGAAAGAATAGTAG, Reverse-outer: ATTTTCATGTACCCGCCCTTGA; Forward-inner: CTACACGACGCTCTTCCGATCT, Reverse-inner: AGACGTGTGCTCTTCCG. The cycling conditions were 95 °C for 3 min; 25 cycles (for outer PCR) and 15 cycles (for inner PCR) of 95 °C for 20 s, 60 °C for 10 s, and 70 °C for 10 s; 70 °C for 5 min. In parallel, the plasmid vector pool was used as a template for the same PCR amplification. The PCR products were indexed and sequenced with paired-end ∼150 bp reads on an Illumina HiSeq, NovaSeq, or MiniSeq instrument.

### Sequencing data analysis

Reads were trimmed for adapters and quality using Trimmomatic^44^ in paired-end mode. Editing outcomes were analyzed using CRISPResso2 (v2.2.13)^22^ using CRISPResso Batch mode with default (Cas9) parameters including the ignore_substitutions option. For the lentiviral integrant sequence analysis, we identified UMIs from the plasmid pool associated with copies of the barcode and candidate sequence perfectly matching the design, aka dud removal. We limited the analysis of sequences amplified from genomic DNA to this set of non-dud UMIs since this procedure reduced the background indel estimate from non-edited cells while maintaining accuracy, precision, and power.

### Editing proportion estimation

We quantified editing activity at each candidate sequence by processing with a custom R script (https://www.R-project.org/) the Alleles_frequency_table_around_sgRNA output file generated by CRISPResso2. We considered the read count as a reliable quantification of the abundance of a specific allele configuration in a sample. Therefore, for each combination of candidate sequence/donor/replicate and condition (edited/unedited), we estimate the total proportion of edited alleles by summing the read counts of alleles having the *“*Unedited” flag set to “FALSE.” We modeled the proportion of *IndelReads* over *TotalReads (*Edited+Unedited) for a given candidate sequence using a binomial response generalized linear model (logistic regression) including the following covariates: *DonorID*, factor distinguishing cell donor; *Treatment,* a binary variable having value *Treatment*=1 for *Edited* and *Treatment*=0 for *Unedited* samples. From the regression analysis results, we calculated the *IndelReads* proportion and the related 95% confidence interval. We considered the editing proportion meaningful when above the technical background noise set to 0.1%, to take into account errors introduced by the PCR amplification steps and the sequencing platform.

## DATA AVAILABILITY

The sequencing data supporting the findings of this study are available on [SRA] under accession number [XXX]. ABSOLVE-seq analytical pipeline is publicly available on GitHub at https://github.com/pellinlab/ABSOLVE-seq. For any inquiries regarding data access or code usage, please contact the corresponding authors.

## ACKNOWLEDGMENTS

L.P. received support from the U.S. NIH (R35 HG010717 and RM1 HG009490). D.E.B. was supported by the National Heart, Lung, and Blood Institute (OT2HL154984, P01HL053749), Doris Duke Charitable Foundation and the St. Jude Children’s Research Hospital Collaborative Research Consortium.

## AUTHOR CONTRIBUTION

Conceptualization, L.P., D.P, D.E.B.; Methodology, S.W., C.B., L.P., D.P, D.E.B.; Investigation, J.L., M.A.N., L.Y.L., J.Z., A.V., N.R.N., L.F.S., A.M., K.L.S., K.C.; Formal analysis, J.L, M.A.N., L.Y.L, D.P., K.C.; Writing – Original Draft, D.P. and D.E.B.; Writing – Review & Editing, L.P, D.P. and D.E.B and all authors; Funding Acquisition, D.E.B; Supervision, L.P, D.P. and D.E.B

## SUPPLEMENTARY FIGURES

**Supplementary Figure 1.**
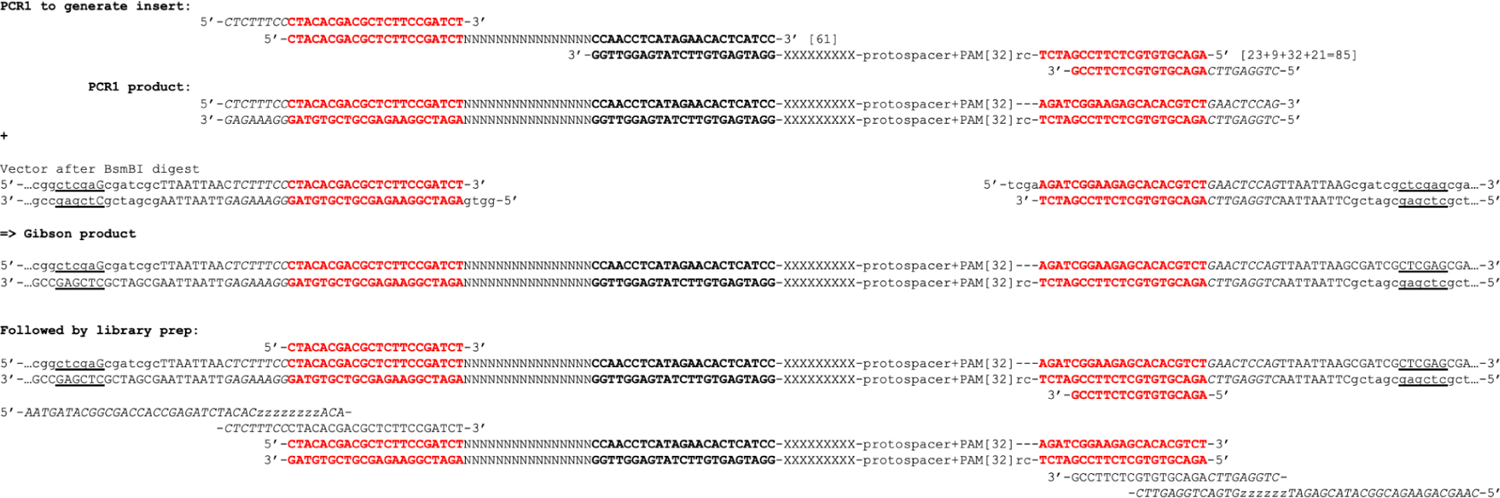
Insert design, cloning strategy, and library preparation. Handles for amplicon sequencing in red, BsmBI recognition sequence in bold, and XhoI recognition sequence in underline. The molecular UMI is denoted by NNNNNNNNNNNNNNNN, candidate off-target barcode by XXXXXXXXX, and sample index by zzzzzzzz. “Protospacer+PAM[32]” entails the 3 nt endogenous upstream sequence, 20 nt protospacer, 3 nt PAM, followed by 6 nt endogenous downstream sequence for a total of 32 nt.

**Supplementary Figure 2.**
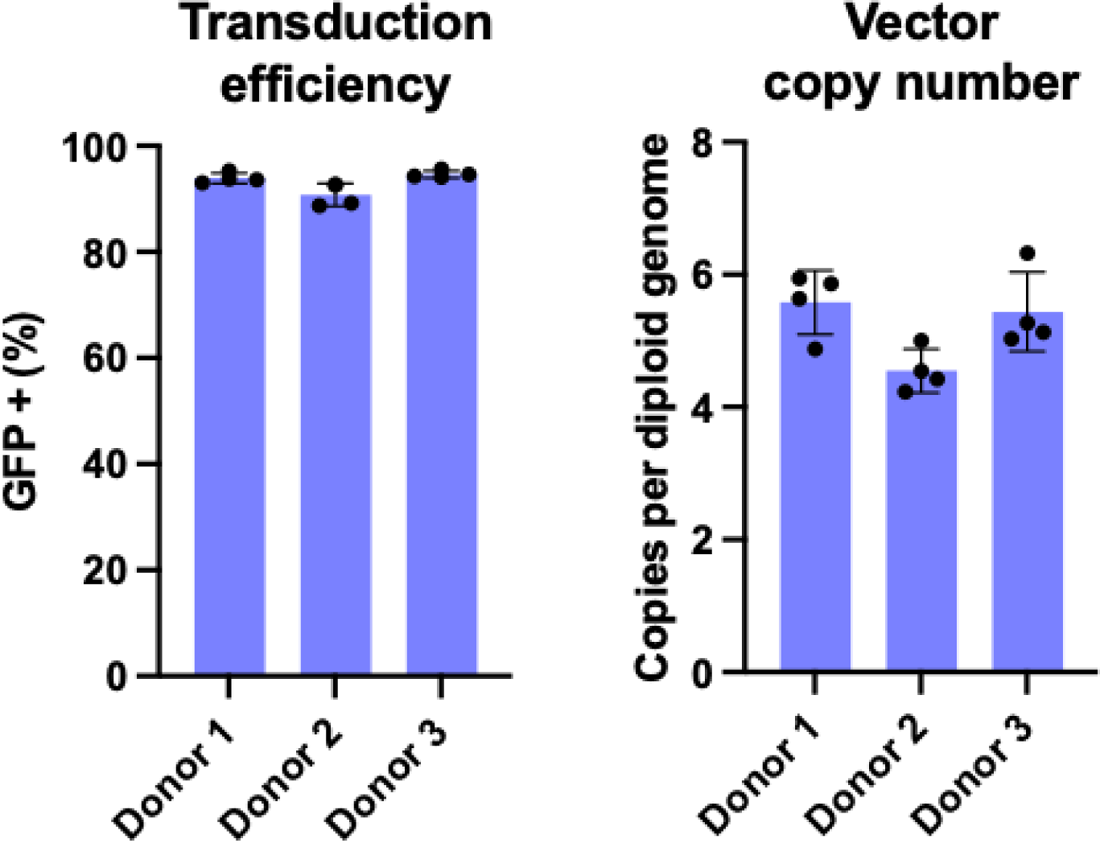
Vector transfer. (A) Transduction efficiency as measured by GFP+ cells using flow cytometry. (B) Vector copy number using ddPCR.

**Supplementary Figure 3.**
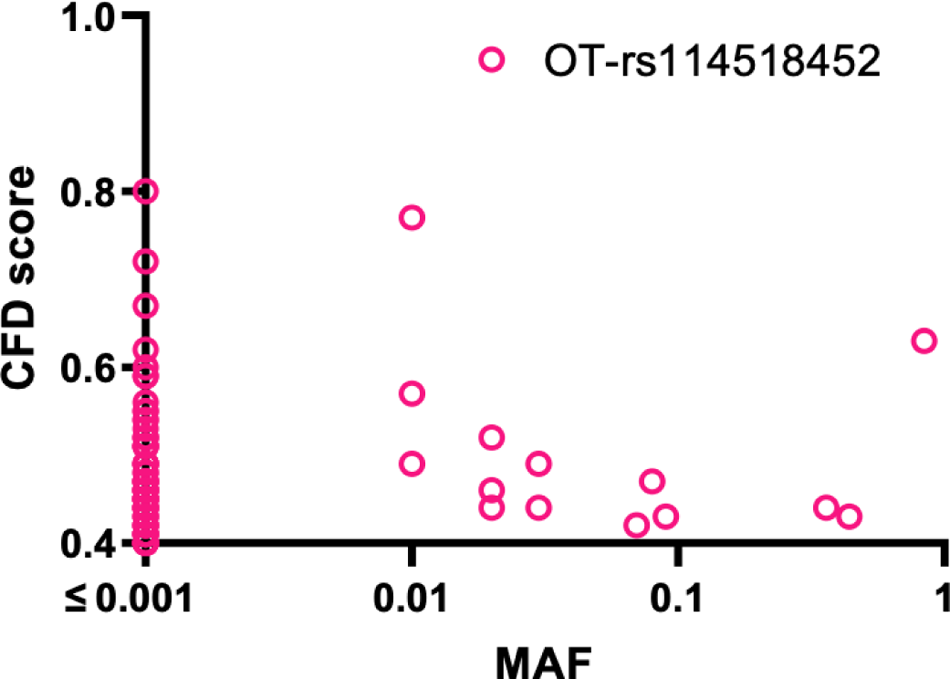
CFD vs Minior Allele Frequency (MAF) for 73 candidate variant-associated off-target with CFD ≥ 0.4

**Supplementary Figure 4.**
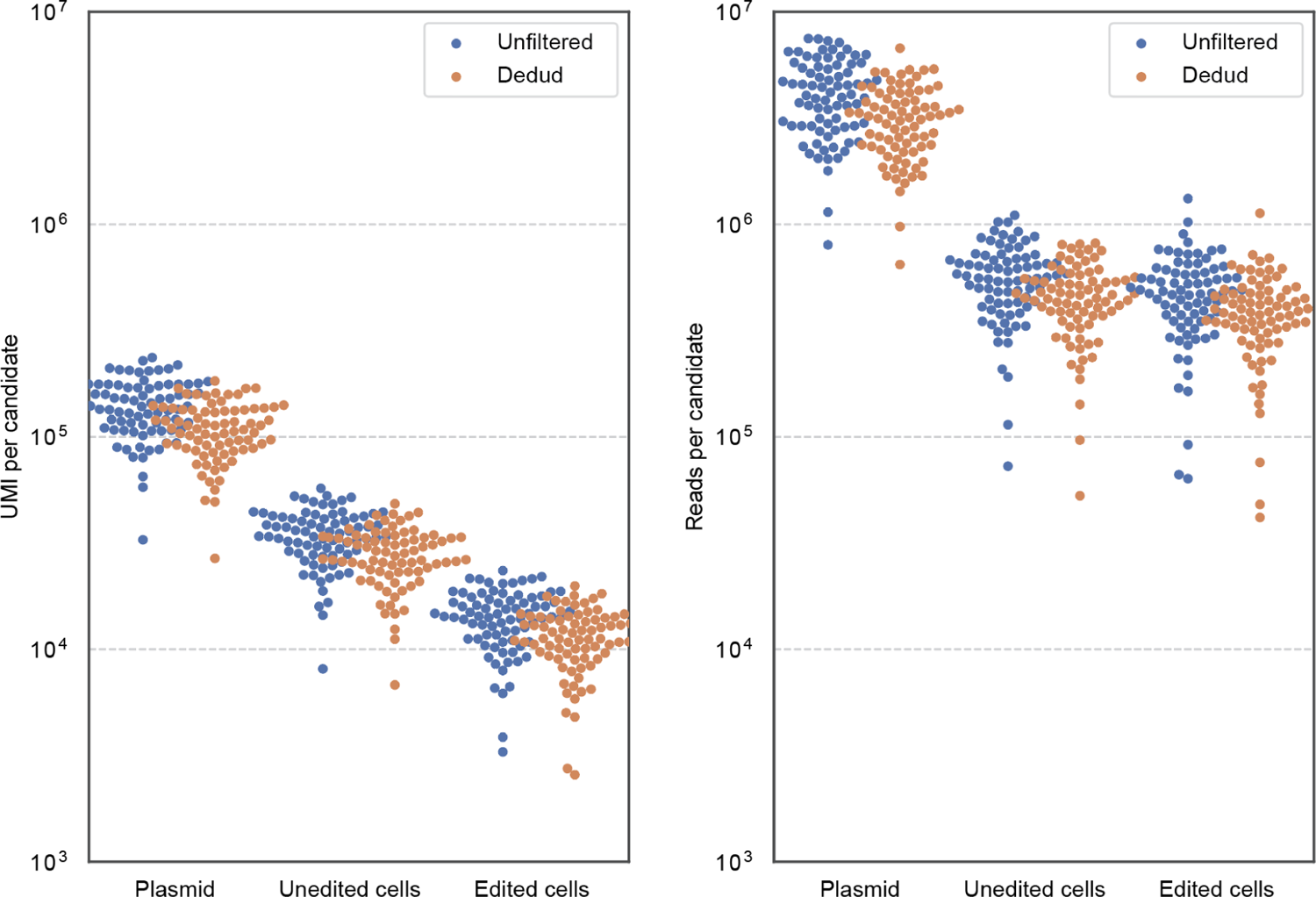
Swarm Plot of UMI and Read Counts for Target Sequences. Swarm plot visualizing the number of UMIs and reads recovered for each target sequence (79) across the plasmid pool, unedited and edited samples. The left panel displays the UMI counts, while the right panel shows the read counts for each target sequence. Each dot represents a target sequence.

**Supplementary Figure 5.**
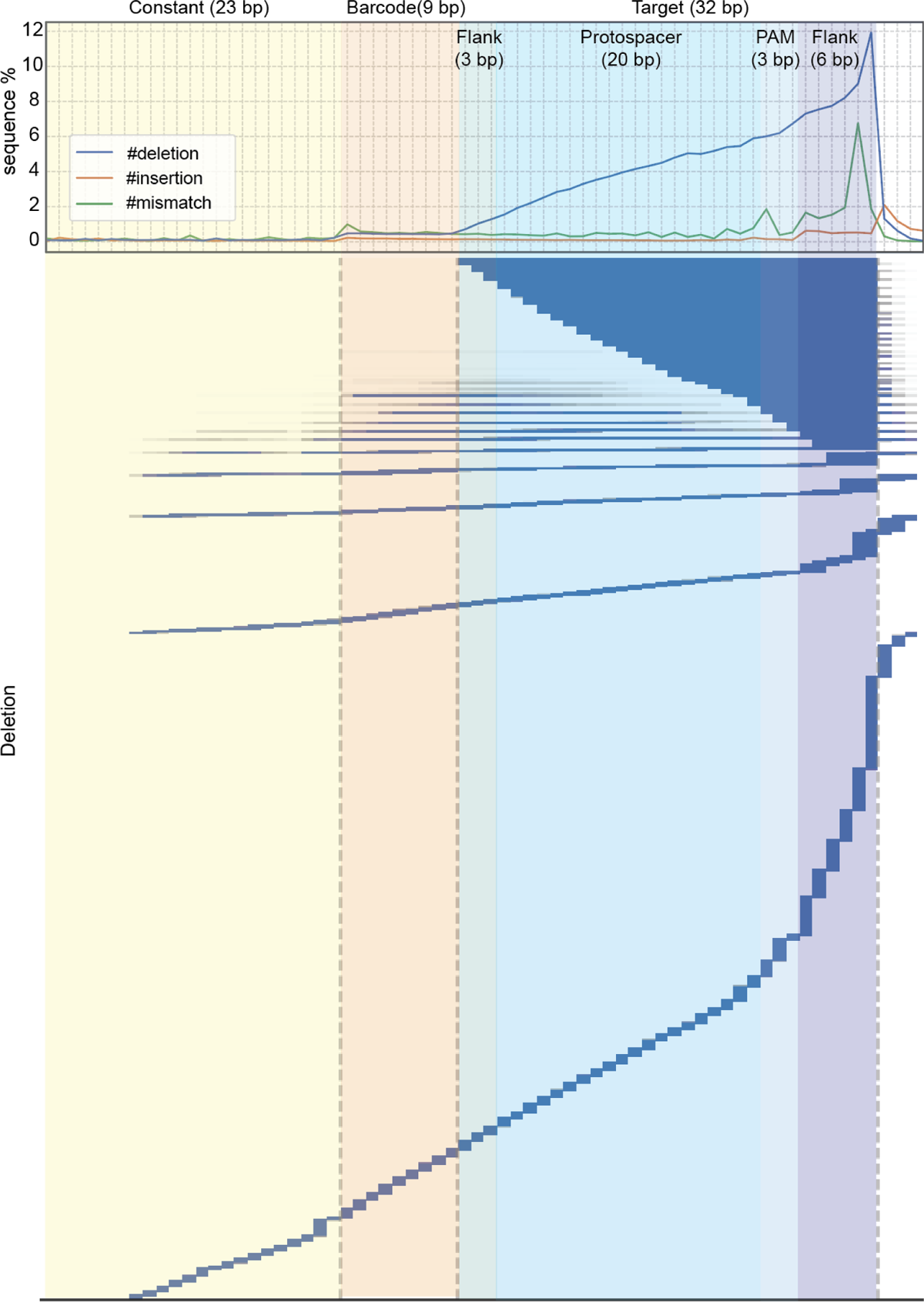
Analysis of the distribution and frequency of deletion, insertion and mismatch in plasmid pool sequencing. Alignment of plasmid pool reads to the expected construct sequence. Top section reports the frequency of deletion, insertion and substitution in the different section of the integrant sequence (constant region, barcode, and target sequence). In the bottom section a detailed representation of abundance and distribution of long (>1 nucleotides) deletion is shown.

**Supplementary Figure 6:**
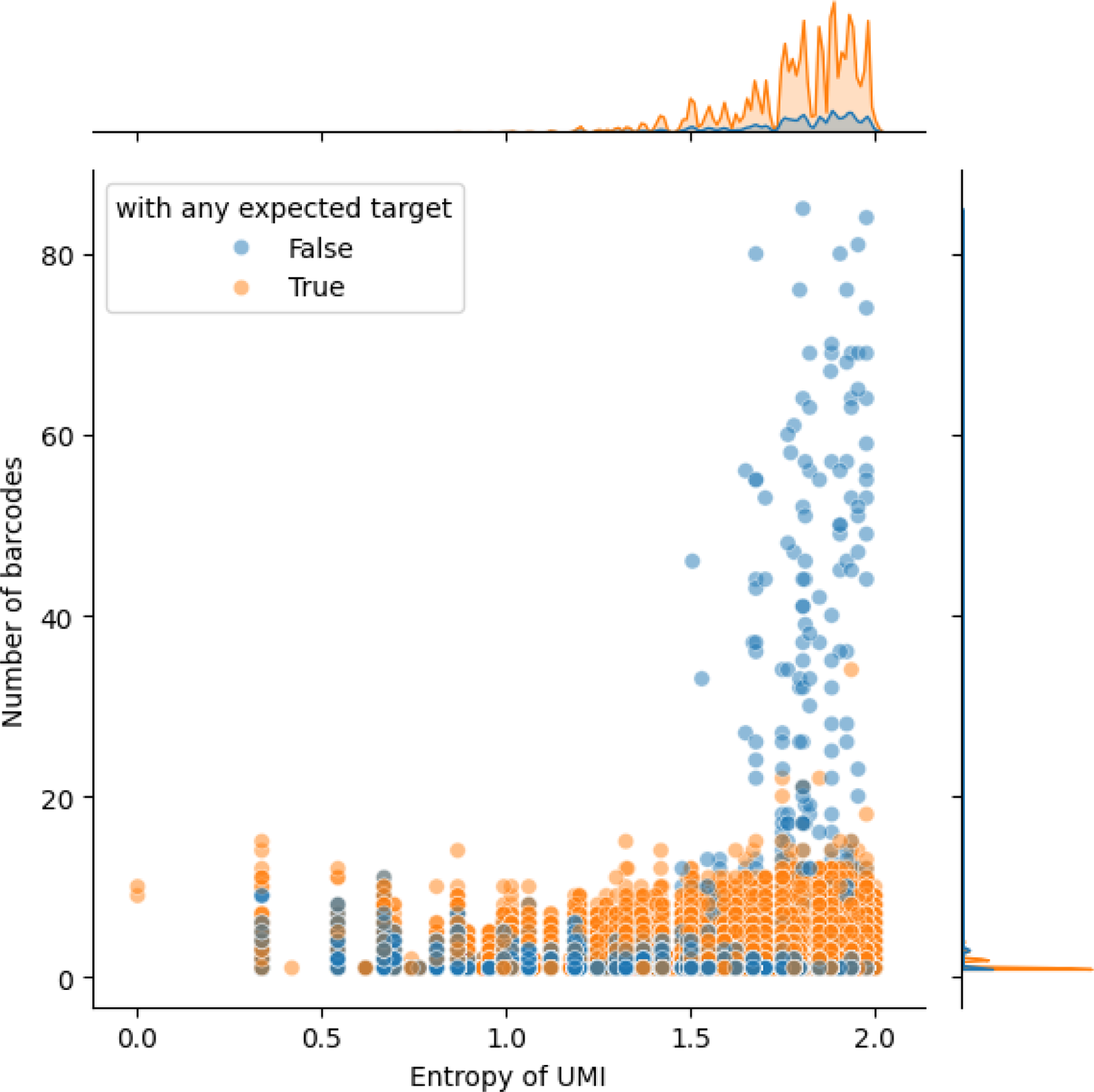
UMI/barcode collision and UMI complexity. Scatter plot illustrating the relationship between UMI complexity and barcode pairing in the plasmid pool. Each dot represents an individual UMI, with orange dots indicating UMIs paired to a candidate barcode and blue dots representing UMIs not paired with any candidate barcode. The x-axis denotes UMI complexity, where a value of 0 indicates a homopolymer UMI. The y-axis shows the number of distinct barcodes that a UMI has been paired with in the plasmid pool.

**Supplementary Figure 7.**
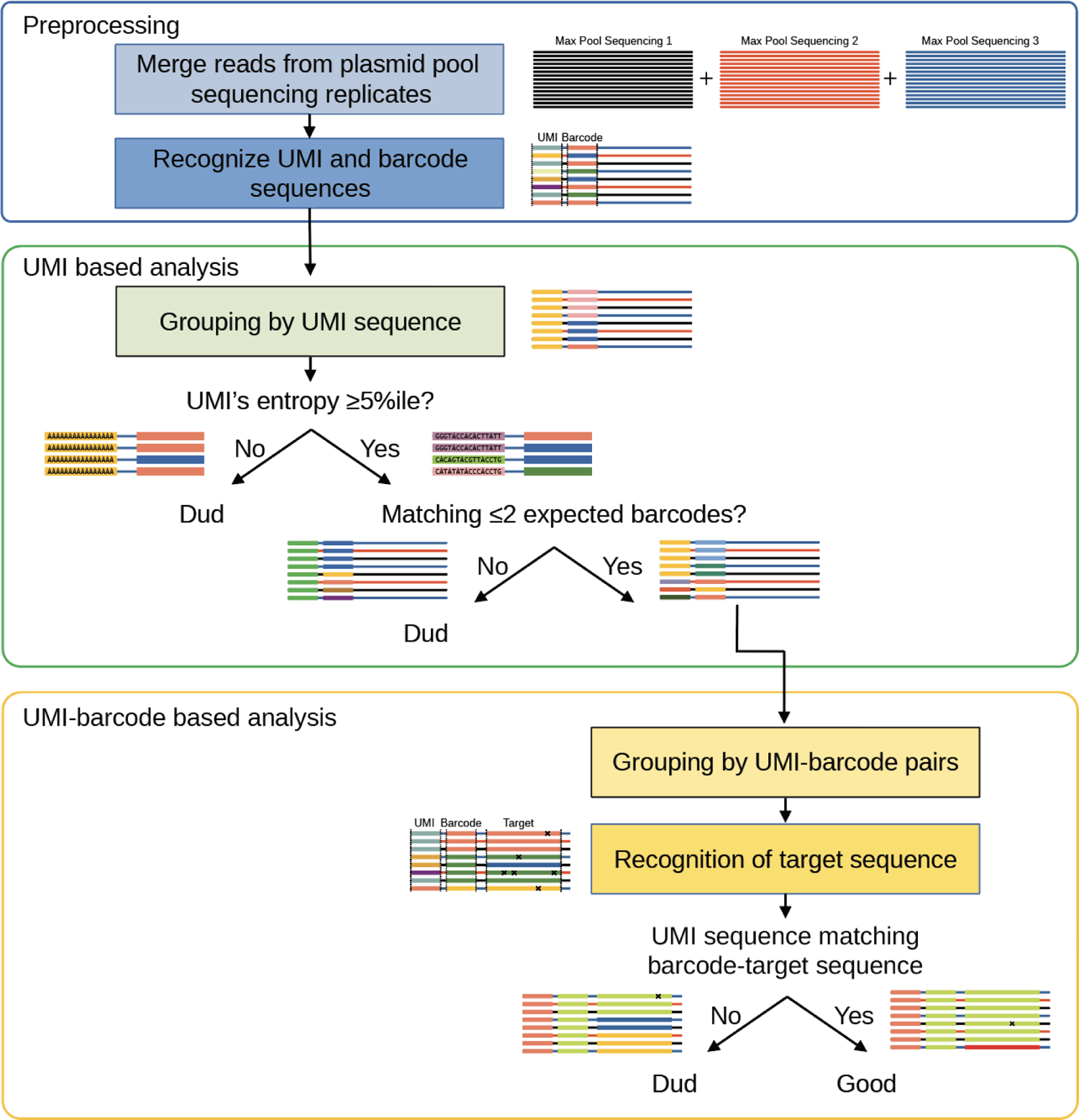
Schematic of Dedud strategy for the identification of high-fidelity UMIs from the plasmid pool sequencing data.

**Supplementary Figure 8.**
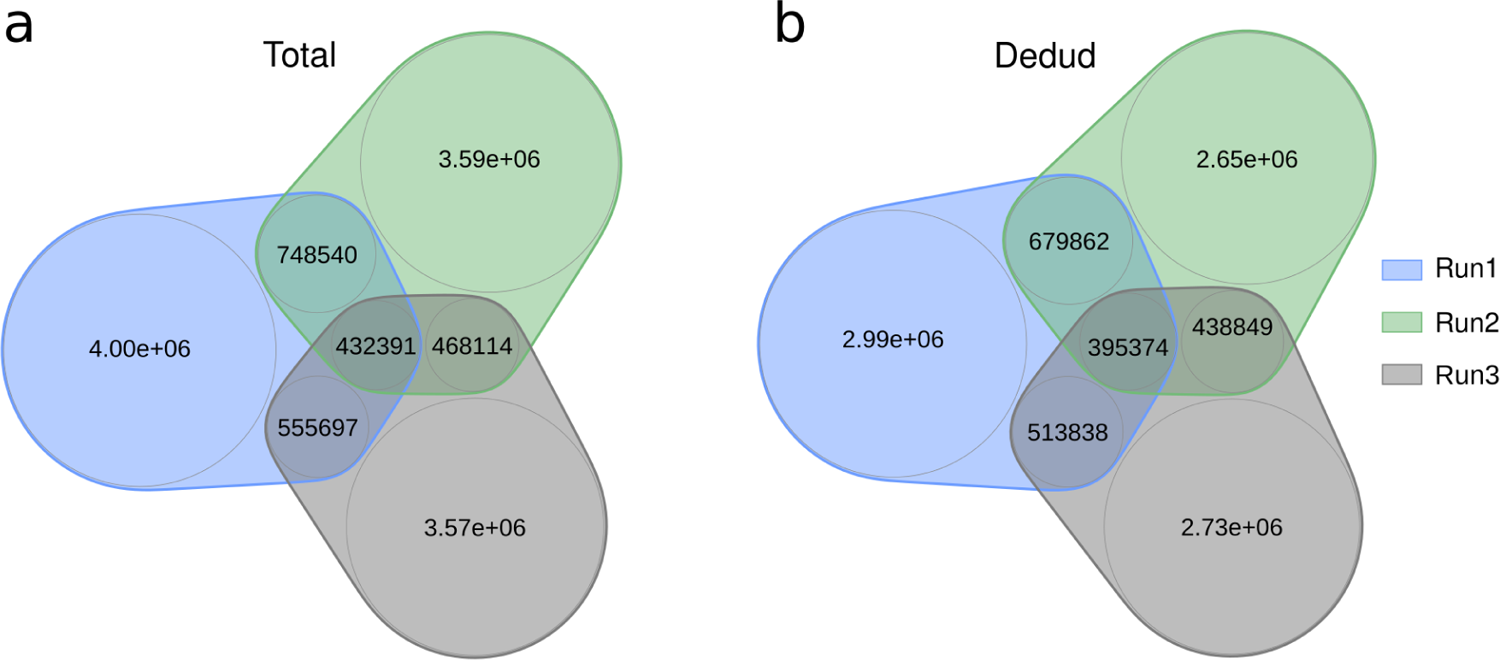
Number of unique and recaptured UMI from three plasmid pool sequencing runs. (A) Total UMIs, (B) UMIs after *Dedud* filtering.

